# Exploration of the Anticancer Activity of *Selaginella bryopteris* derived silver nanoparticles

**DOI:** 10.1101/2025.01.25.634850

**Authors:** Ankita Dutta, Khushbu Wadhwa, Neha Kumari, Chetan Pandey, Anoop Kumar, Neha Kapoor

## Abstract

Ovarian cancer is characterized by chronic hormonal imbalances and associated symptoms like irregular menstruation, fertility problems and abdominal discomfort. Poly ADP-ribose polymerases (PARPs) are proteins that play a crucial role in DNA repair pathways like base excision repair (BER). PARP family proteins exhibit catalytic activities which facilitate the transport of ADP-ribose to targeted proteins. Inhibition of PARP proteins in cancer cells with *BRAC1/2* gene mutations impairs the process of Single Strand Break (SSBs) repairs followed by Double Strand Breaks (DSBs), stalling replication, leading to apoptosis. PARP1 protein inhibition is an innovative therapeutic approach in cancer treatment. Commercially available drugs like Olaparib and Rucaparib are used as PARP1 inhibitors to treat ovarian cancer, however the development of effective new therapeutic drugs is urgently needed to overcome drug toxicity and resistance. In this study, we synthesized silver nanoparticles (AgNPs) from the leaf extract of *Selaginella bryopteris* demonstrating their cytotoxic nature against the ovarian cancer SKOV-3 cell line, with an IC_50_ value of 7.81µg/ml. We also observed cell death in cancer cells with an ROS upsurge induced by *S. bryopteris* AgNPs. Our findings also elaborate on the role of Rhamnetin, a *S. bryopteris* phytochemical that exhibits active site targeting of PARP-1 residues.

## 1. Introduction

Gynaecological cancers, which are one of the most prevalent cancers affecting women globally, can disrupt the normal functioning of the female reproductive system, leading to a considerable decrease in the lifespan. The epidemiological trends of these cancers vary among different geographic regions and change over time (**Kevyani et al., 2023**). In the year 2018, ovarian cancer was identified as the seventh most common cancer in women across the world (**Shafabakhsh et al., 2019**). In India, ovarian cancer is next in importance to cervical cancer and is defined as the most common cancer of the female genital tract. By definition, a malignant tumour in one or both ovaries in females is known as ovarian cancer. All ovarian cell types exhibit the potential to develop into ovarian cancer (**Berek et al., 2018; Kroeger et al., 2017**). The three primary types of ovarian cancer are germ cell tumours, sex-cord stroma tumours and epithelial ovarian cancer (∼90%). The primary cause of ovarian cancer is the existence of mutations in genes associated with the MAPK (mitogen activated protein kinase) pathways ((CDKN2A (Cyclin dependent kinase inhibitor 2A), p-catenin genes, PIK3CA (phosphatidylinositol-4,5-biphosphate 3-kinase, catalytic alpha), TP53 (Tumor protein 53), BRAF (v-Rafmurine sarcoma viral oncogene homolog B)). In addition to this, ovarian cancer is also characterized by suppression of the tumor by inactivating mutations in the *ARIDIA* gene (AT-rich interaction domain 1A) (**Rojas et al., 2016**). Moreover, *ERB2* gene amplification that codes for the HER-2 (human epidermal growth factor receptor 2) is also seen in patients suffering from ovarian cancer. The mode of spread of ovarian cancer is distinct from other cancers due to the extremely rare occurrence of hematogenous metastasis. It spreads by passive diffusion mechanisms, where the cancerous cells are shed off from the tumor’s surface and get transferred by the physiological movement of peritoneal fluid. The accumulation of pathologic peritoneum fluid also leads to the formation of ascites that further facilitate cancer cell spread (**Yeung et al., 2015**). For the management of ovarian cancer, different forms of treatments are available including surgery, chemotherapy, radiation, hormonal and photo therapy (**Summer et al., 2024a**). Chemotherapy, which is widely used in treating cancer has adverse effects involving harmful effects on healthy cells. Additionally, patients undergoing chemotherapy may develop multiple side-effects and drug resistance with high recurrence risks (**Yin et al., 2022**). Chemotherapeutic agents rapidly kill dividing cells; however, majority of the available drugs are unable to distinguish between cancerous and non-cancerous cells. Patients undergoing chemotherapy suffer adverse effects because of modulation of other targets and deleterious effect on normal cells (**Pearce et al., 2017**).

Despite great improvement in these conventional therapies, they still exhibit limitations. For eg., such therapies due to the inability of the drugs involved in penetrating the malignant tissue cause inflammation as a potential side-effect (**Summer et al., 2024a**).

In contrast to this, targeted therapies are directed towards cancer specific mutations and inhibit tumor growth. This minimizes the negative effect on the nearby non-malignant cells (**Torino et al., 2013; Anand et al., 2022**). Targeted therapies not only give more positive outcomes but also result in fewer off-target side effects. Thus, it is very essential to develop drugs that overcome challenges exhibited by conventional therapies and only target cancerous cells (**Summer et al., 2024a**).

In recent years therapeutic and diagnostic approaches based on nanotechnology have also shown the ability to improve cancer therapy (**Ratan et al., 2020**). Nanotechnology has received attention due to distinctive properties exhibiting broad-range applications in various fields including agriculture, electronics, textile industry, cosmetics, medicine, environment remediation and healthcare settings (**Mohammad et al., 2022**). Among the various metallic nanoparticles available, silver nanoparticles stand out as a highly commercialized choice worldwide because of their physicochemical and biological characteristics (**Summer et al., 2024b**). Nanoparticles usually range in size from 1 to 100 nm and exhibit unique physical and chemical properties. The biocompatibility of nanoparticles depends on their size and as their size decreases, their surface reactivity increases. Smaller sized nanoparticles exhibit more ability in being absorbed by cells causing minimal damage (**Summer et al., 2024c**).

Silver nanoparticles (AgNPs) can be synthesized by employing diverse methods including chemical reduction, physical vapor deposition, electrochemical methods, and green or biological synthesis (**Iravani et al., 2014**). They can be obtained by synthesis with biological agents like bacteria, fungi, algae and various plant parts. Biological synthesis of silver nanoparticles has several advantages with lesser production of toxic products, enhanced biocompatibility, reduced toxicity towards healthy cells, low cost, easy maintenance and rapid production (**Mandal et al., 2006; Gericke and Pinches, 2006**).

The green synthesis of silver nanoparticles from plant extracts minimizes the use of hazardous chemical agents and make them biocompatible and ecofriendly in nature as compared to conventionally used chemical processes (**Zulfiqar et al., 2024**). Abbas et al., 2024 reported the radical scavenging activity of Zinc oxide nanoparticles synthesized from the leaves, flowers and bud extracts of *Bauhinia variegata* indicating the role present phytocompounds like polyphenol, tannins, alkaloids, amides, saponins, glycosides and flavonoids in capping and stabilization of these nanoparticles.

Researchers have found nanoparticles to exhibit minimal potential risks, ensuring their safe usage (**Mishra et al., 2015**) and significant potential as anticancer and antiangiogenic agents (**Choi et al., 2016**). In this study, we synthesized AgNPs using aqueous leaf extracts of *Selaginella bryopteris* with silver nitrate.

Our previous published work aimed at characterising *S. bryopteris* obtained AgNPs (**Wadhwa et al., 2023a**). Many phytochemicals including curcumin, quercetin, resveratrol, silymarin that are derived from medicinal plants are generally used to treat different types of cancers by inhibiting the process of angiogenesis and induction of apoptotic cell death pathways. In a recent study, researchers demonsrated the anticancer activity of silymarin loaded Ni-Fe metal-organic frameworks against breast cancer and neuroblastoma cancer cell lines (**Rahimi et al., 2024**). In addition to this, a nonregulator was assembled through coordination bonds between phytic acid and Fe^3+^ to synthesize nano-sized metal-organic frameworks (MOFs), with mitoxantrone (MTO) as a loaded drug that was tested against metastatic bone tumors (**Huang et al., 2023**). Natural products are considered as major source of anti-cancer compounds as they have ability to inhibit the proliferation of various types of cancers. One of the sesquiterpenes, ‘elemene’ isolated from the rhizome of *Curcuma wenyujin*, is linked with anti-tumor pharmacophore named nitric oxide that forms β-elemene NO donor hybrids. These hybrids are further tested for anti-cancer activities against lung squamous cell carcinoma cell lines (NCL-H520), glioblastoma cell line (U87MG) and colorectal cell lines (SW620) (**Bai et al., 2022**). Xanthohumol, a prenylated flavonoid isolated from a species of flowering plant, *Humulus lupulus* (common name ‘hops’), has exhibited anticancer activity against gastric cancer cell lines. The decreased cell viability was attributed to inhibition of proliferation and induction of apoptosis upon treatment with xanthohumol (**Wei et al., 2018**). Yadav et al., 2020 reported the antibacterial activity of *S. bryopteris* derived silver nanoparticles against *M. luteus*, *P. mirabilis*, *B. megaterium* and *E. coli* by disc diffusion and agar well diffusion methods.

### 1.1 Selaginella bryopteris

*Selaginella bryopteris* (L.) Bak or Resurrection plant is a desiccation tolerant medicinal plant belonging to the Selaginellaceae family (**Pandey et al., 2010**). It is also known as “Sanjeevani booti” (that infuses life) for its therapeutic and medicinal properties. The crucial role of this herb has been even listed in great historical epics like *Ramayana*. *S. bryopteris,* which is a perennial, herbaceous and lithophytic plant, grows on rocks and is mostly found in humid environments. It has special leaves called “microphylls” along with distinct structures known as ligules, which are outgrowths near the upper surface base. Rhizophores bear the roots and the spore bearing structures, sporangia, which are present in the axils of sporophylls (**Antony and Thomas, 2011**). During the summer season, this plant undergoes extreme desiccation, and its fronds become curled and dried. When monsoon arrives, *Selaginella* turns green and comes back to its active life (**Sah et al., 2005**). The plant is also an example of unspecialized primitive heterospory (**Schulz et al., 2010**) and plays a key role in evolutionary studies showing potent adaptation to xeric conditions (**Figure 1**).

**Figure 1:**
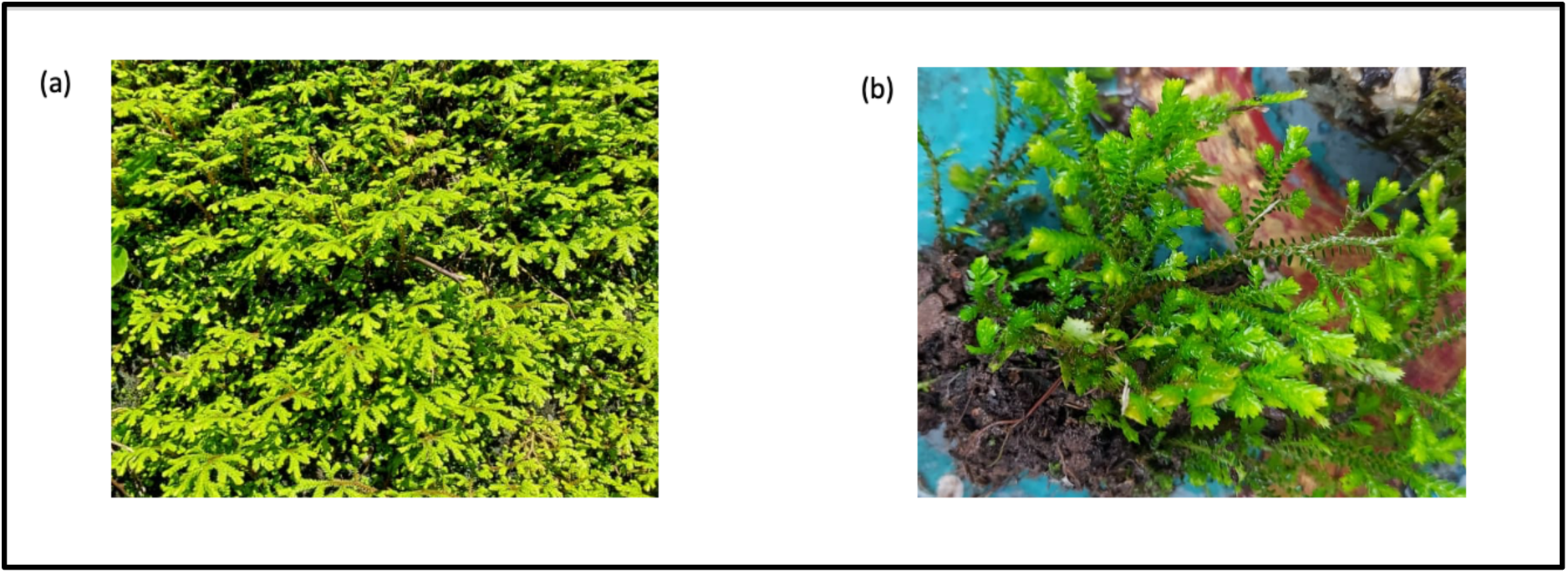
(a) *Selaginella bryopteris* in its natural habitat (b) Plant morphology

*Selaginella* contains several types of secondary metabolites including flavonoids, phenolics, and alkaloids that play a vital role in defence mechanisms. Secondary metabolites from the plant act as reducing and stabilizing agents for nanoparticle synthesis. The plant has been used since ancient times for treating fever, kidney problems, asthma, and diarrhoea *S. bryopteris* ethanolic extracts have been found to exhibit significant wound healing properties (**Paswan et al., 2020**). GC-MS analysis has indicated the presence of flavonoids, bioflavonoids, sugars, sugar alcohols, glycosides, campesterol, stigmasterol and β-sitosterol (**Paswan et al., 2020; Gautam et al., 2023**).

### 1.2 PARP1 and associated Single-Strand Break Repair (SSBR)

The group of proteins called poly (ADP-ribose) polymerase (PARP) family has attracted much attention for cancer treatment in recent years. They are also known as ADP-ribosyl transferase diphtheria-toxin-like proteins (ARTD1–17). PARP proteins contain two ribose sugars and two phosphate moieties per unit polymer. In humans, there are 17 PARPs, each containing common catalytic domains composed of ∼230 amino acids (**Lu et al., 2019**). Even though they have "poly-ADP-ribose" in their names, only four of these enzymes (PARP1, 2, 5a, and 5b) are the ones that create long chains of poly-ADP-ribose and attach them to their target molecules (**Burke and Virag, 2013**). PARP proteins are crucial in having significant roles in DNA repair, recombination, transcription, DNA replication (**Figure 2 A**) and modulation of chromatin structure, ensuring proper function of centromeres and formation of mitotic spindles. They also support centrosome function, control telomere dynamics, facilitate the movement of endosomal vesicles, and affect cell death through apoptosis (**Burkle, 2005**).

**Figure 2:**
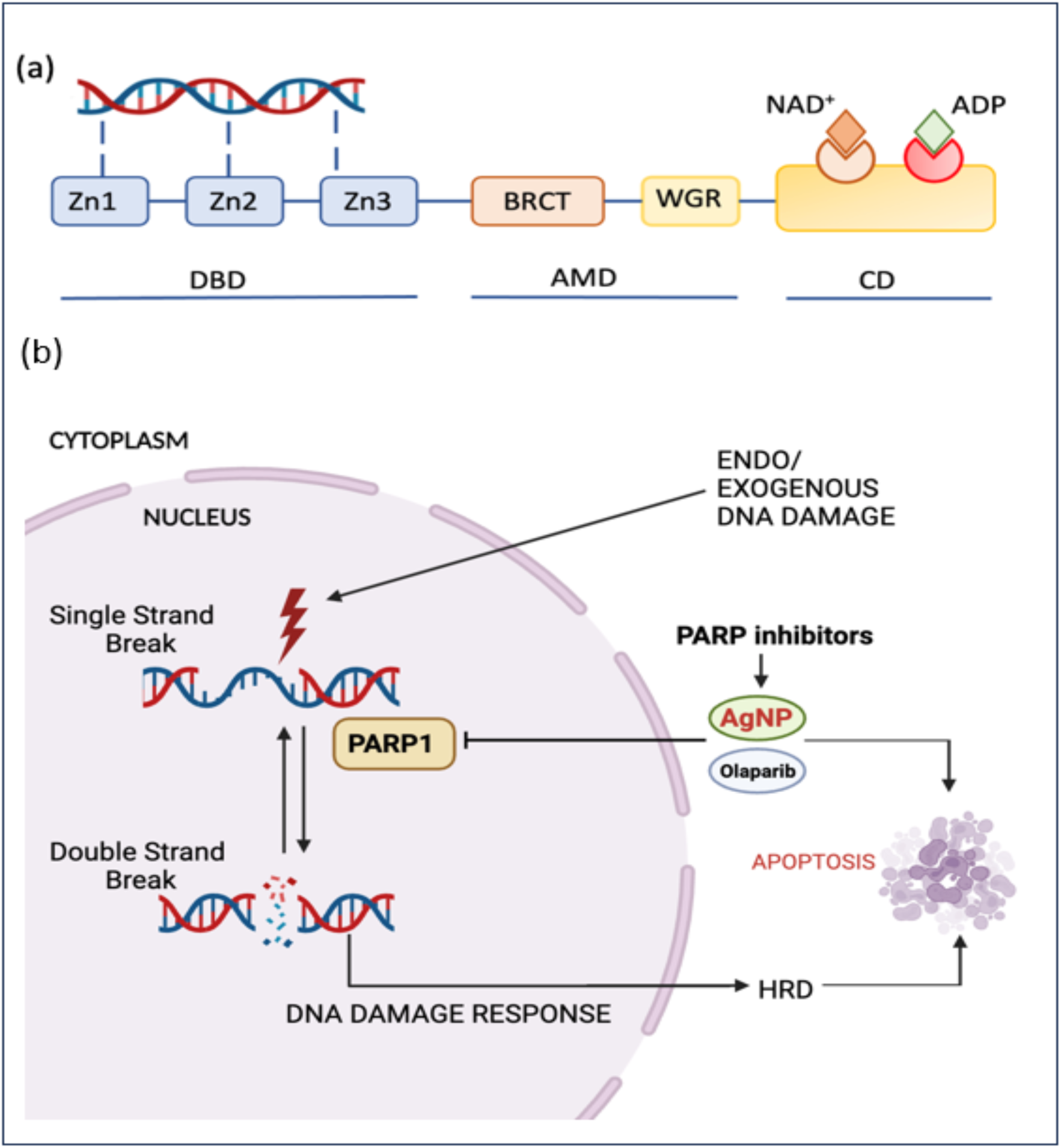
(a) Schematic representation of various functional domains of PARP-1 protein including DBD (DNA binding domain), AMD (Auto-modification domain), CD (Catalytic domain). (b) Proposed mode of action of PARP1 inhibitors (AgNPs, Olaparib) upon exo/endogenous DNA damage in cancer cells.

PARP protein is an important protein in DNA repair pathways of base excision repair (BER) (**Chen and Du, 2018**). BER is known to preserve and maintain the integrity of DNA during cellular oxidative stress in response to any exogeneous damage induced by toxic chemicals or mutations. It is involved in DNA repair during single strand breaks (SSBs) and if due to some mutations BER is impaired or PARP is inhibited, it causes accumulation of SSBs leading to occurrence of double strand breaks (DSBs). Homologous recombination (HR) and non-homologous end joining (NHEJ) mechanisms can be altered due to impairment of BER and PARP in the cells. It was found that patients whose cells are defective in HR and NHEJ are even more susceptible to impairment of BER pathways that can lead to progression of cancer (**Maynard et al., 2009**).

PARP1 and PARP5A make up most of the PARP family members that are major subjects of research for scientists (**Kleine et al., 2008**). The most common protein of this family is PARP-1, which is found in the nucleus. If there is DNA damage, it is majorly PARP-1 which is involved in repair (**Rojo et al., 2012**). PARP-1 is an ADP ribosyl transferase superfamily enzyme which has DNA dependent activity and catalyzes the synthesis of poly (ADP ribose) from NAD^+^ molecules. The binding of PARP-1 protein to DNA breaks results in the reorganization of chromatin structure and recruitment of DNA repair proteins to repair damage. For designing competitive inhibitors of PARP-1, detailed understanding of the molecular interactions of proteins and substrates is a primary requirement. PARP-1 protein contains three zinc DNA binding domains at the N-terminal region that helps this protein to recognize breaks in DNA. The second domain, referred to as the auto-modification domain (AMD), has a tryptophan-glycine-arginine (WGR)-rich domain and a centrally regulating section that contains a motif from the C terminus of the breast cancer susceptibility protein (BRCT) (**Figure 2 b**) (**Nilov et al., 2020; Langelier and Pascal, 2013**) The catalytic domain of PARP-1 protein has two binding sites named as donor (NAD^+^) binding site and the acceptor (PAR) binding site (**Figure 2 b**). The NAD^+^ molecule binds to PARP protein by forming two hydrogen bonds with Gly 863 and hydrophobic contacts (pi-stacking) with Tyr 907. With this, adenosine diphosphate (ADP) forms a hydrophobic contact with Met 890 along with hydrogen bonds with two residues named as Lys 903 and Glu 988 (**Nilov et al., 2020**).

### 1.1 Necessity of PARP Inhibitors

Various PARP (Poly ADP-ribose polymerase) enzyme inhibitors were approved and available in the market for the treatment of several types of cancers, particularly in the context of targeting BRCA (breast cancer gene) mutations. These inhibitors are primarily required for the treatment of curing breast, ovarian, and other cancers with DNA repair deficiencies. The commercially available inhibitors are Niraparib, Rucaparib, Olaparib, Talazoparib and Veliparib. Although, these inhibitors are commercially available in the market, but there is a pressing need to develop novel PARP inhibitors because these drugs exhibit a large number of adverse effects, including fatigue, elevated cholesterol levels, headaches, dizziness, nausea, vomiting, renal toxicities, including elevated creatinine concentrations (**LaFargue et al., 2019**).

## 2. Results

### 2.1 Silver Nanoparticles and Their Characterization

#### 2.1.1 Biosynthesis of AgNPs

A significant color change in the solution preliminarily confirms the green synthesis of silver nanoparticles. The color changes from pale brown to dark brown, indicating the reduction of silver ions in the aqueous solution by the capping agent present in the leaf extract of *Selaginella bryopteris* (**Figure 3**).

**Figure 3:**
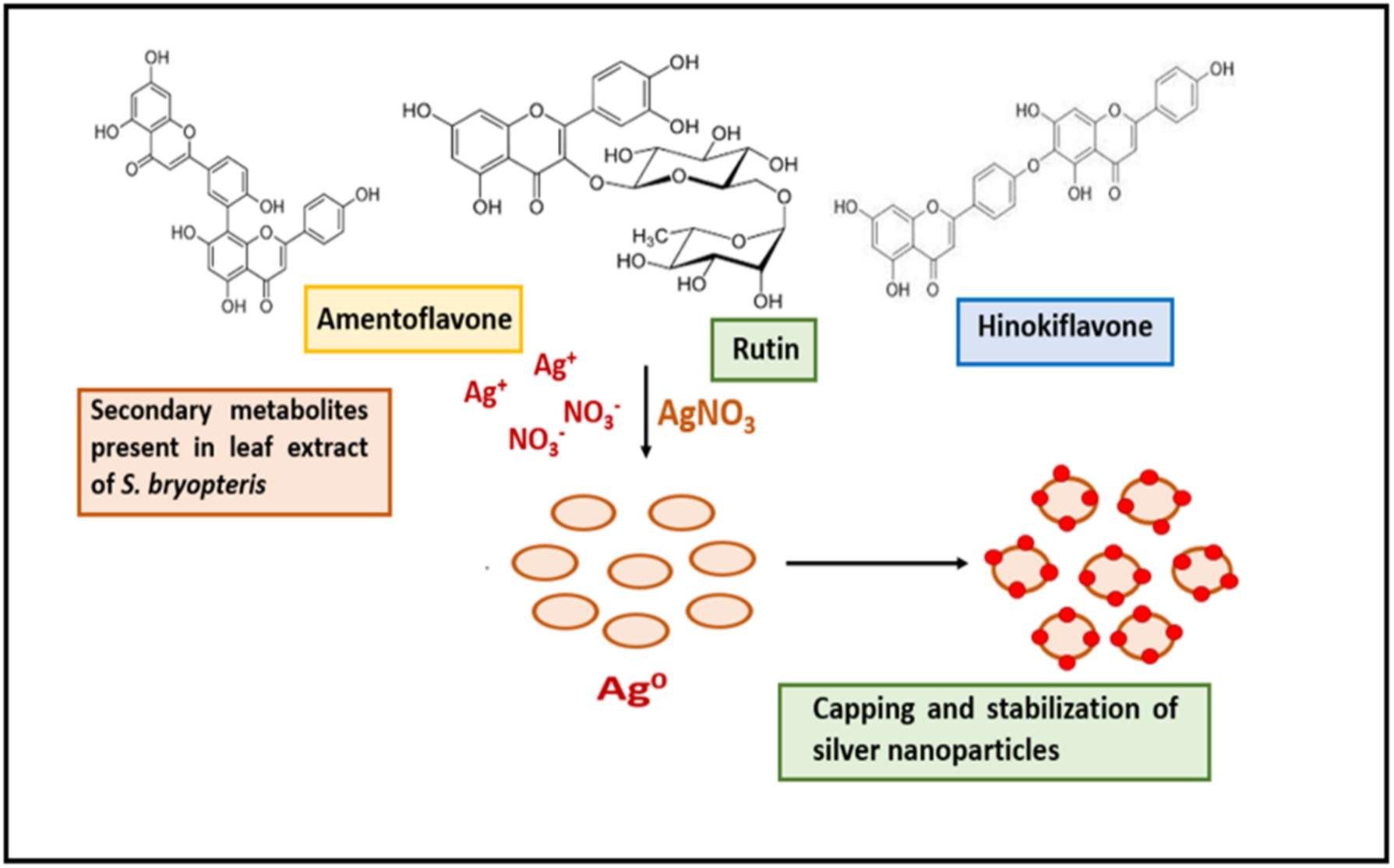
Schematic representation of mechanism of plant mediated biosynthesis of silver nanoparticles using aqueous leaf extract of *S. bryopteris*.

#### 2.1.2 Characterization of green synthesized silver nanoparticles

In one of our previous studies (**Yadav et al., 2020**), *S. bryopteris* derived silver-nanoparticles have been characterized by FTIR and X-Ray diffraction (XRD) techniques as shown in (**Figure 4**).

**Figure 4:**
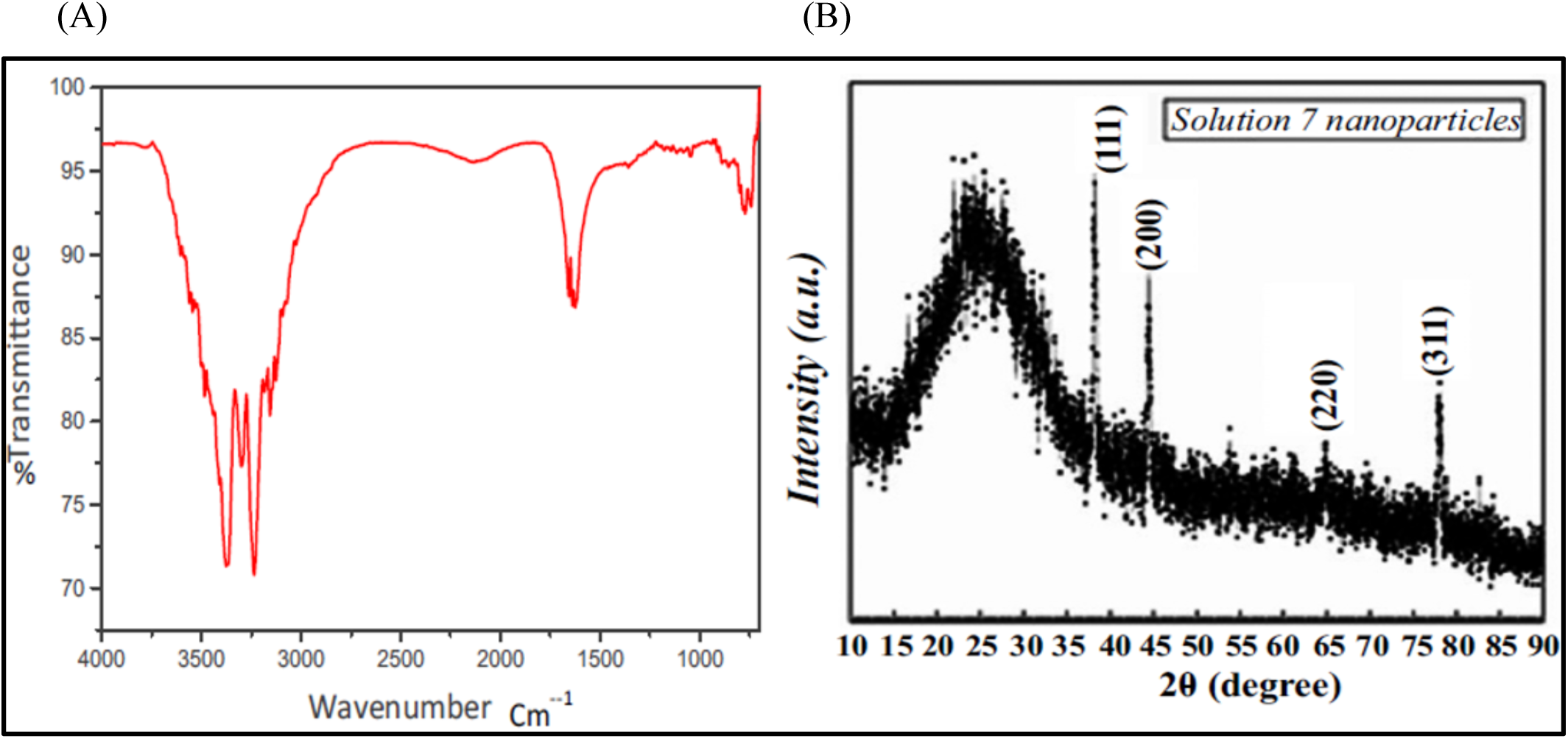
- (A) FTIR spectra and (B) XRD pattern of green synthesized silver nanoparticles using leaf extract of *S. bryopteris*.

The analysis of the FTIR spectra revealed prominent absorbance bands at approximately 1046, 1356, 1626, 1656, 2132, 3156, 3236, 3300, 3376, and 3484 cm^-1^ indicating various functional groups and chemical bonds present at the surface of silver nanoparticles as confirmed by previous studies done by Mughal et al., 2024; Tahir et al., 2024; Tanveer et al., 2024. The observed peak at 1046 cm^-1^ indicates a C-C stretch, C-N aromatic amino acid or C=C bond of an aromatic ring. Bands at 1356 cm^-1^ indicate C-C and C-N stretching whereas at 1626 cm^-1^ represent the stretching vibrations of primary amines. Bands at 3156, 3236, 3300 and 3376 cm^-1^ indicate the presence of phenol and alcohol groups and a peak at 3484 cm^-1^ indicating O-H stretching vibrations of phenol group (**Figure 4 A**). Thus, the FTIR spectra confirms that biomolecules present in the leaf extract of *S. bryopteris* such as flavonoids, bioflavonoids and alkaloids helped in capping and stabilization of silver nanoparticles. XRD analysis shows four distinct diffraction peaks at (2ϴ) - 38.22°, 44.62° and 64.96° and 77.52° corresponding to (111), (200), (220) and (311) planes of silver (**Figure 4 B**). FESEM images show that green synthesized silver nanoparticles exhibit a spherical shape. The elemental analysis of AgNPs was confirmed by energy dispersive X-ray analysis (EDX), and a strong signal of the peak was observed at 3 KeV, which corresponds to absorption of metallic silver nanoparticles as shown in **Figure 5**. Furthermore, the shape size and morphology of AgNPs was confirmed by transmission electron microscopy (TEM). The TEM image shown in **Figure 6** clearly indicates that AgNPs are spherical in shape. (**Figure 6**)

**Figure 5.**
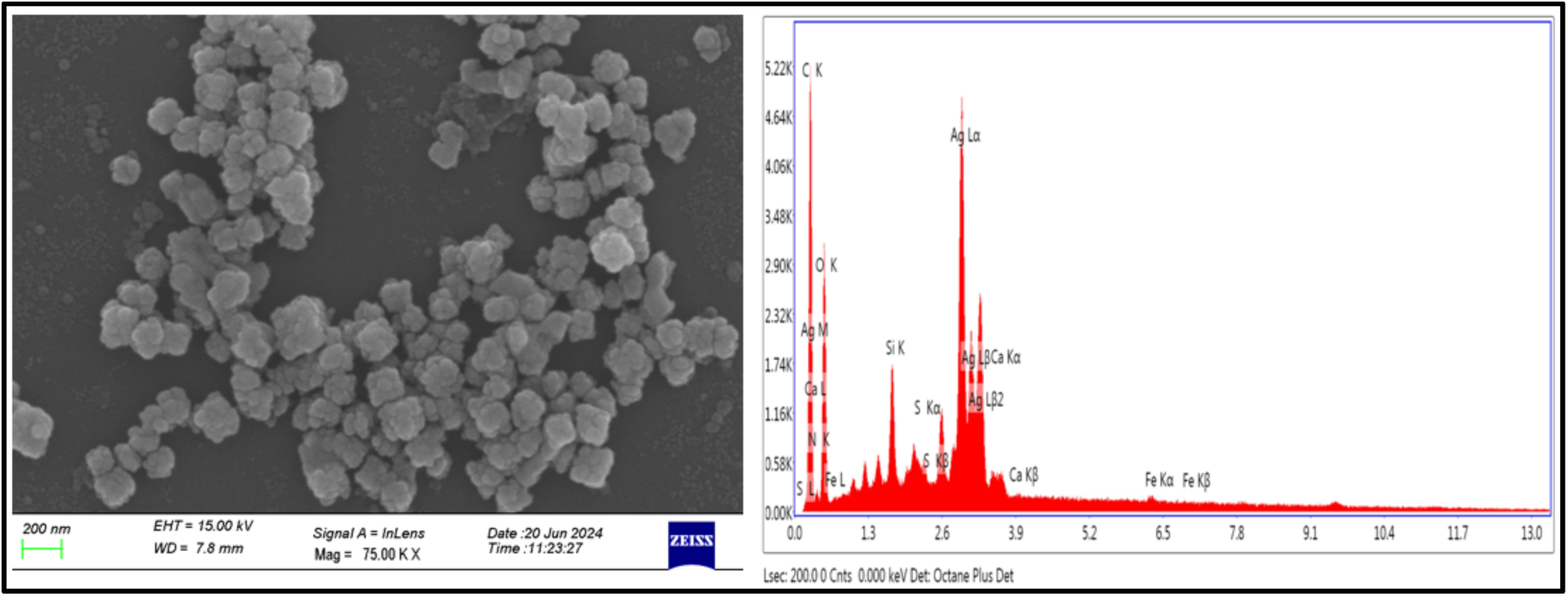
FESEM and EDX mapping of green synthesized AgNPs using aqueous leaf extract of *S. bryopteris*.

**Figure 6:**
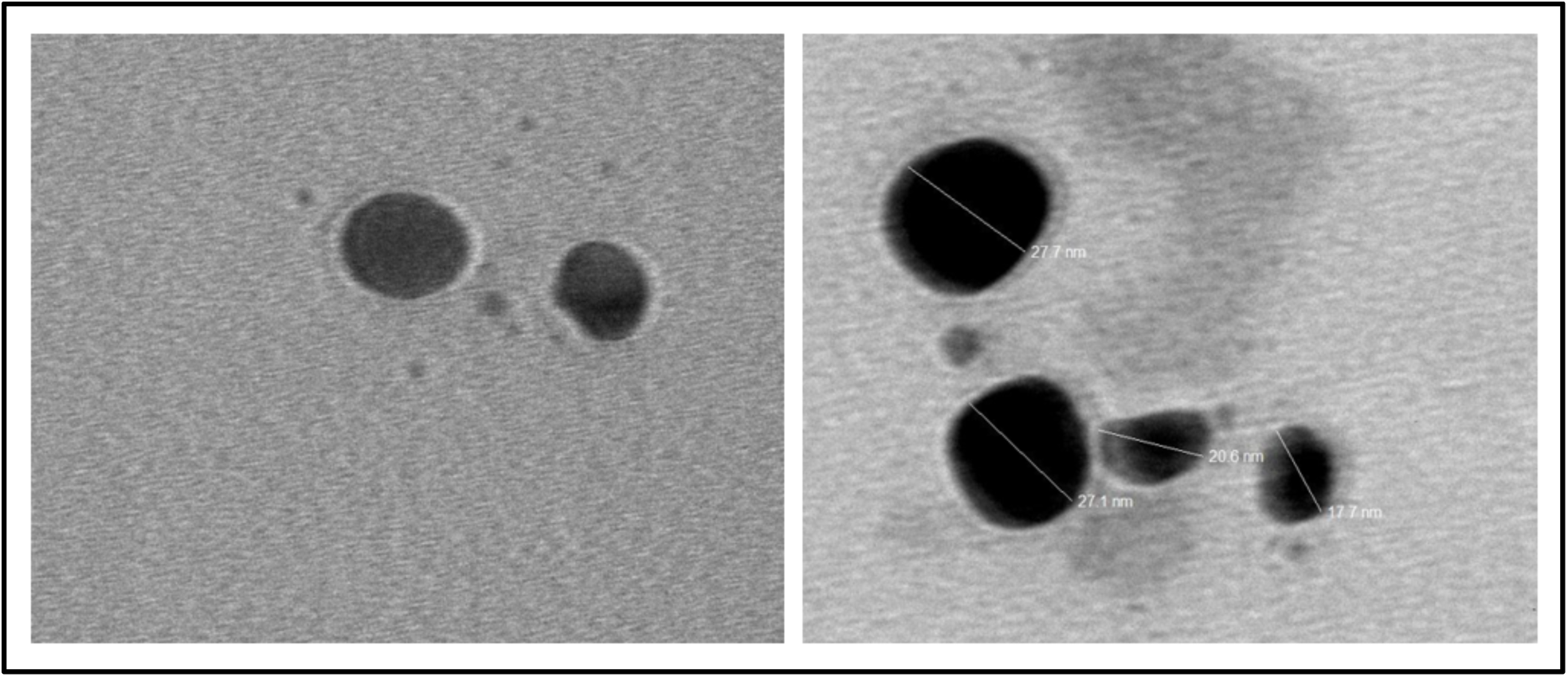
TEM images of spherical biosynthesized AgNPs.

The ICP-MS analysis helps to analyze different inorganic components or micronutrients present in the leaf extract of medicinal *S. bryopteris* plant (**Table 1**). Trace elements play very crucial role in the formation of chemical constituents of plants. Some metals such as zinc, iron, chromium, copper and cobalt are not highly toxic to some concentrations but as their concentration increases, they become toxic in nature, with this, some other metals including mercury, cadmium, lead are more toxic even at their low concentrations. Thus, it is very necessary to have knowledge about the chemical composition of plant extracts, to formulate our medicines to a safer extent. (**Table 1**)

**Table 1.** Identification of elements from the dried leaf extract of *S. bryopteris* (concentration expressed in ppb-parts per billion)

| Name of element | Mass | Concentration | Units |
| --- | --- | --- | --- |
| Na | 23 | 63.483 | ppb |
| Zn | 66 | <0.000 | ppb |
| Se | 77 | 11.099 | ppb |
| Se | 78 | 7.709 | ppb |
| Se | 82 | 6.817 | ppb |
| Li | 7 | 7.298 | ppb |
| Na | 23 | 53.224 | ppb |
| Mg | 24 | 48.893 | ppb |
| Al | 27 | 18.775 | ppb |
| K | 39 | 96.593 | ppb |
| Ca | 43 | 19.175 | ppb |
| Ca | 44 | 64.880 | ppb |
| Cr | 52 | 0.700 | ppb |
| Mn | 55 | 2.956 | ppb |
| Fe | 56 | 18.597 | ppb |
| Co | 59 | 0.592 | ppb |
| Ni | 60 | 1.215 | ppb |
| Cu | 63 | 0.787 | ppb |
| Zn | 66 | <0.000 | ppb |
| Se | 78 | 5.954 | ppb |
| Sr | 88 | 1.134 | ppb |
| Te | 125 | 30.593 | ppb |
| Ba | 137 | 0.922 | ppb |
| Tl | 205 | 28.201 | ppb |
| [Pb] | 206 | 3.131 | ppb |
| [Pb] | 207 | 3.065 | ppb |
| Pb | 208 | 3.068 | ppb |
| Bi | 209 | 4.455 | ppb |

#### 2.1.3. pH and thermal stability of green synthesized silver nanoparticles

pH is a crucial factor that facilitates the stabilizing and coating ability of phytocompounds. It plays a crucial role in determining the particle size of nanoparticles as it can modify the functional groups of phytocompounds which further alter their reducing and capping ability (**Akhtar et al., 2024a**). The pure stock AgNPs exhibited a clean and narrow SPR with a peak absorbance of 2.5 at 423 nm. Addition of 10% HNO_3_, lowered the peak absorbance to 1.75 and 1.25 with a very broadened SPR. The peak obtained for pH solution 3 and 5 indicated a broad peak of absorption spectra indicating non-uniform (heterogenous) silver nanoparticles. At pH (3 and 5), the SPR broadens indicating the formation of larger sized particles that were formed due to increased particle aggregation and at pH 7, 8 and 9, maximum formation of silver nanoparticles is observed in comparison to stock with a peak absorbance of 3.0, 3.4 and 3.75, SPR at 423 nm. (**Figure 7 A, B**). From the UV-VIS results, basic pH has a better effect on stability of AgNPs as compared to acidic pH (**Figure 7 A**). The AgNPs have higher absorbance at higher pH values (**Akhtar et al., 2024a; Subhani et al., 2024**). The effect of temperature on the stability of AgNPs was evaluated at 3, 55, 70°C and at room temperature. It was found that maximum absorption SPR peak was obtained at 70°C and it remains stable due to the maximum kinetic energy exhibited by AgNPs. The increase in absorbance of silver nanoparticles corresponds to enhanced rate of synthesis of nanoparticles (Figure 8 A and B). (**Akhtar et al., 2024b**). (**Figure 8**)

**Figure 7.**
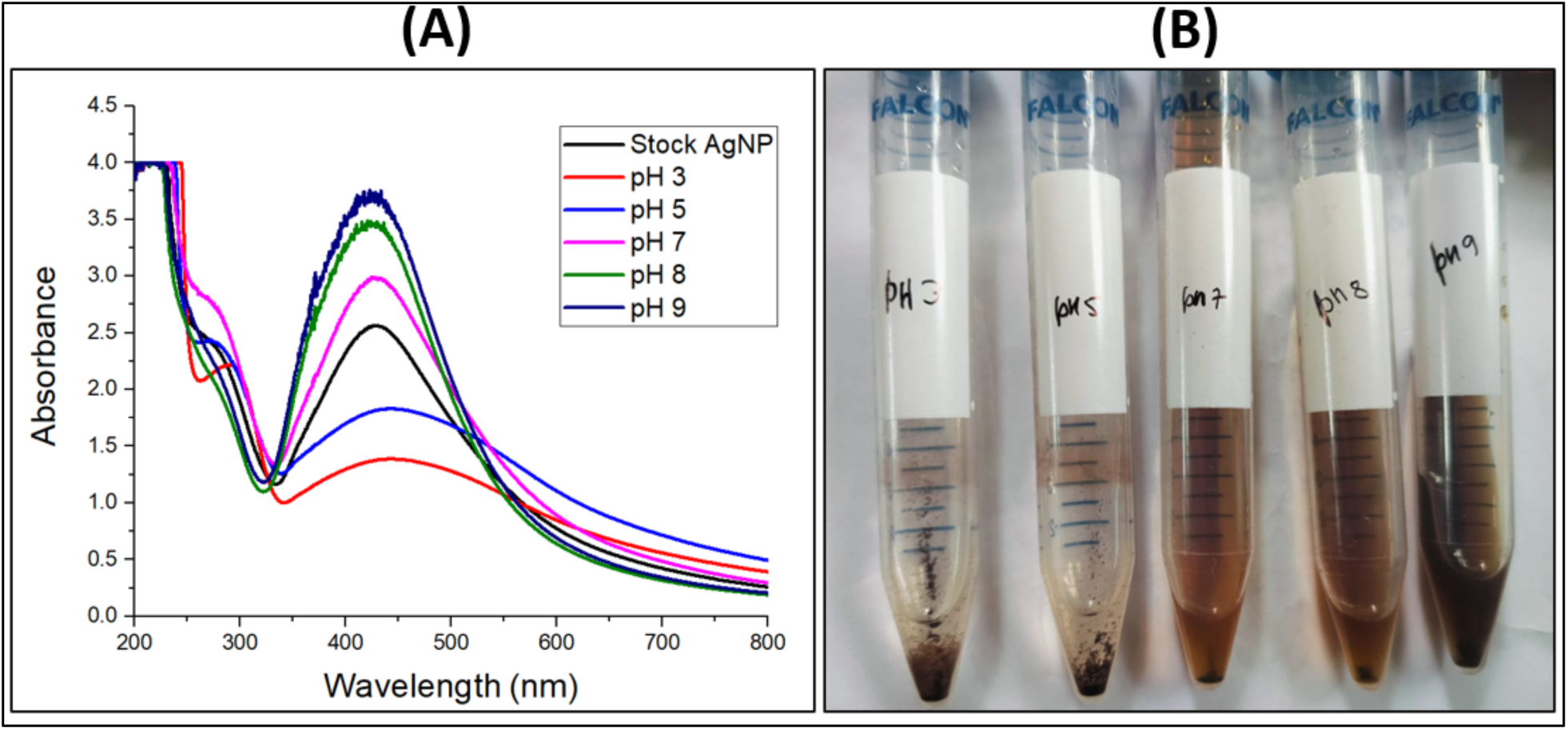
(A) UV-visible spectra of green-synthesized AgNPs at different pH, (B) Visual appearance of AgNPs at different pH.

**Figure 8.**
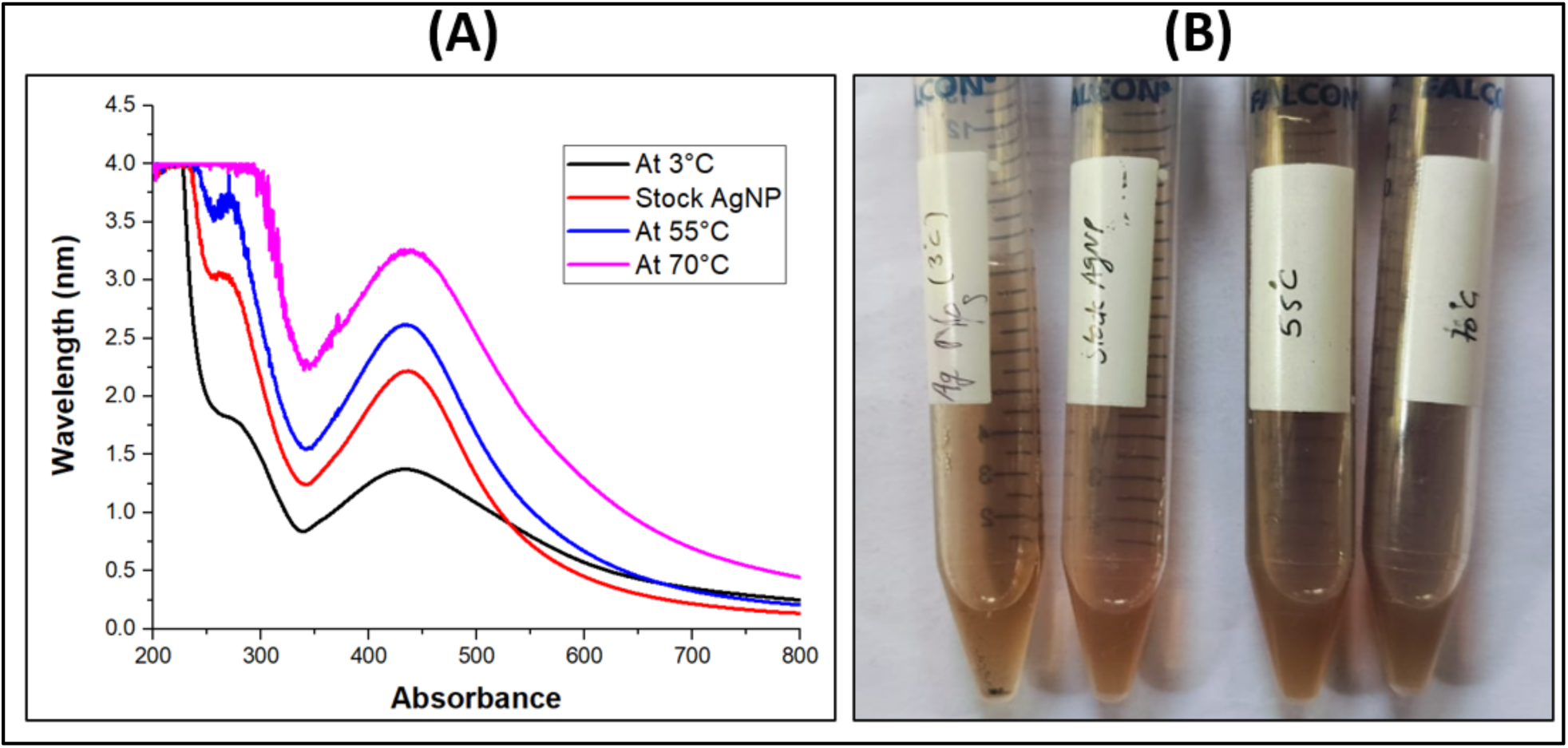
(A) UV-visible spectra (B) Visual appearance of AgNPs at different temperatures to investigate stability of green-synthesized AgNPs.

#### 2.1.4 MTT Assay

MTT assay is used to evaluate the viability of cells receiving a pharmacological treatment. Well-known cytotoxic medications target and prevent cell division by interfering with normal physiological processes. Our results indicate that silver nanoparticles, which are concentration-dependent and decrease cell viability as the drug concentration rises, are the primary source of toxicity. Lower concentrations of the drug (ranging between 200-400 µg/ml) did not exhibit any detectable toxicity. This may be because of the cells’ self defence mechanisms or ineffectiveness of a lower concentration. Significant cytotoxicity was observed starting from a concentration of 500µg/ml. The drug concentration required for reducing the cell viability to its initial 50% (IC_50_) was also calculated from the equation depicted in the cytotoxicity graph (**Figure 9**).

**Figure 9:**
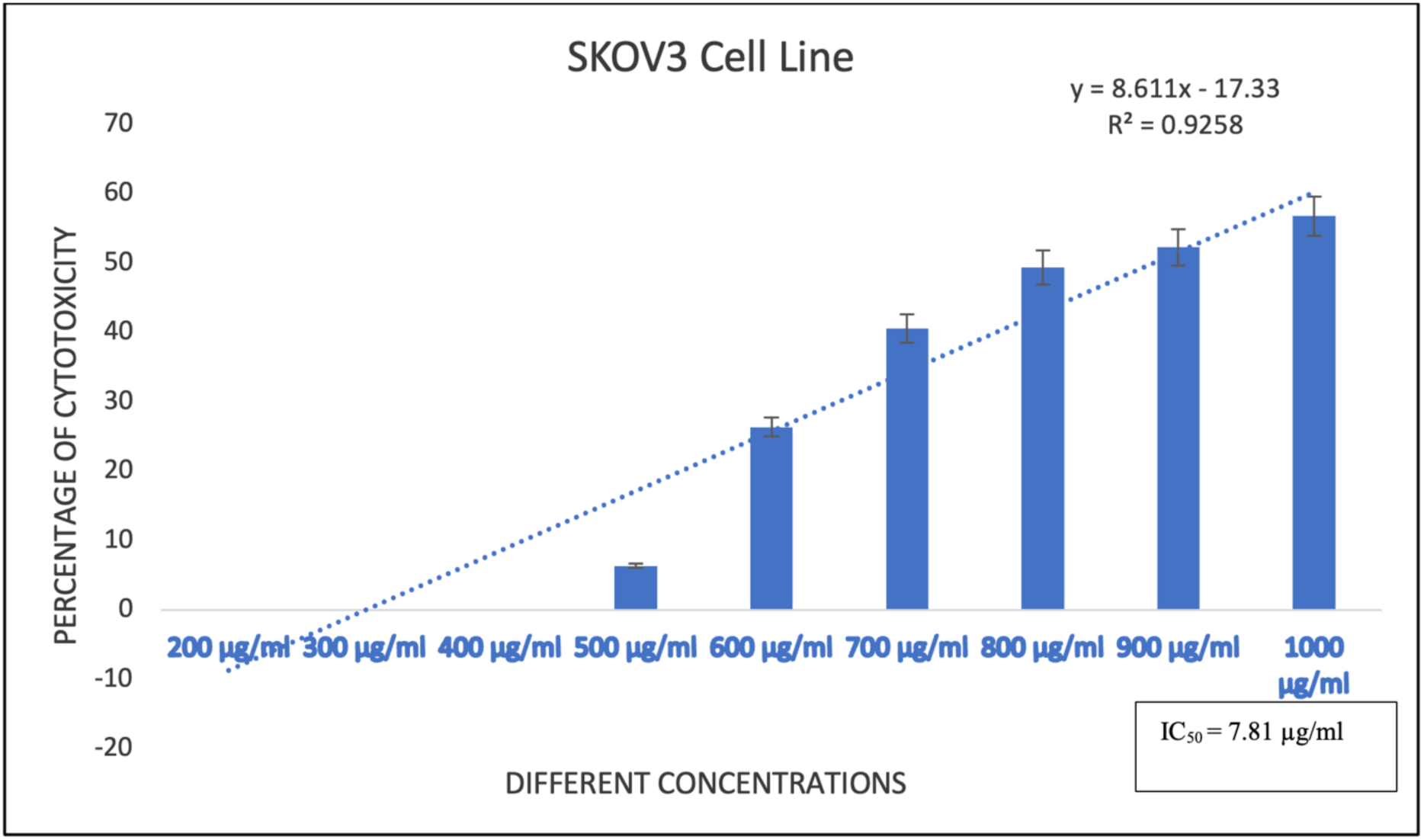
Graph showing the cytotoxic percentage versus different concentration of silver nanoparticles using MTT assay analysis to determine the IC_50_ value of silver nanoparticles against SKOV3 cell line.

This discovery implies that silver nanoparticles, whose activities are concentration-dependent, cause reduction in cancer cell viability. Reactive oxidizing molecules which can be radicals and non-radicals (H_2_O_2_) have hazardous effects on living cells. They can be exogenously or endogenously produced. These molecules can easily take away electrons from (oxidize) molecules with which they remain in contact similar to cellular biomolecules. This initial reaction can generate chain reactions that lead to apoptosis. We have determined the internal ROS production by 2՛,7՛-dichloro dihydrofluorescein diacetate (H_2_DCFA) (non-fluorescein in reduced form) as an indicator for reactive oxygen species (ROS) in the cells. The nonfluorescent H_2_DCFDA converts to significantly fluorescent 2՛,7՛-dichlorofluorescein (DCF) upon cleavage of acetate groups by intracellular esterases and oxidation which could be detected by fluorescent intensity. We have found very minute fluorescence in the control and ROS induced fluorescence in the positive control. The production of ROS was also high in the presence of silver nanoparticles which indicates that ROS can be the possible reason of induction of apoptosis in the cells treated with silver nanoparticles (**Figure 10**).

**Figure 10:**
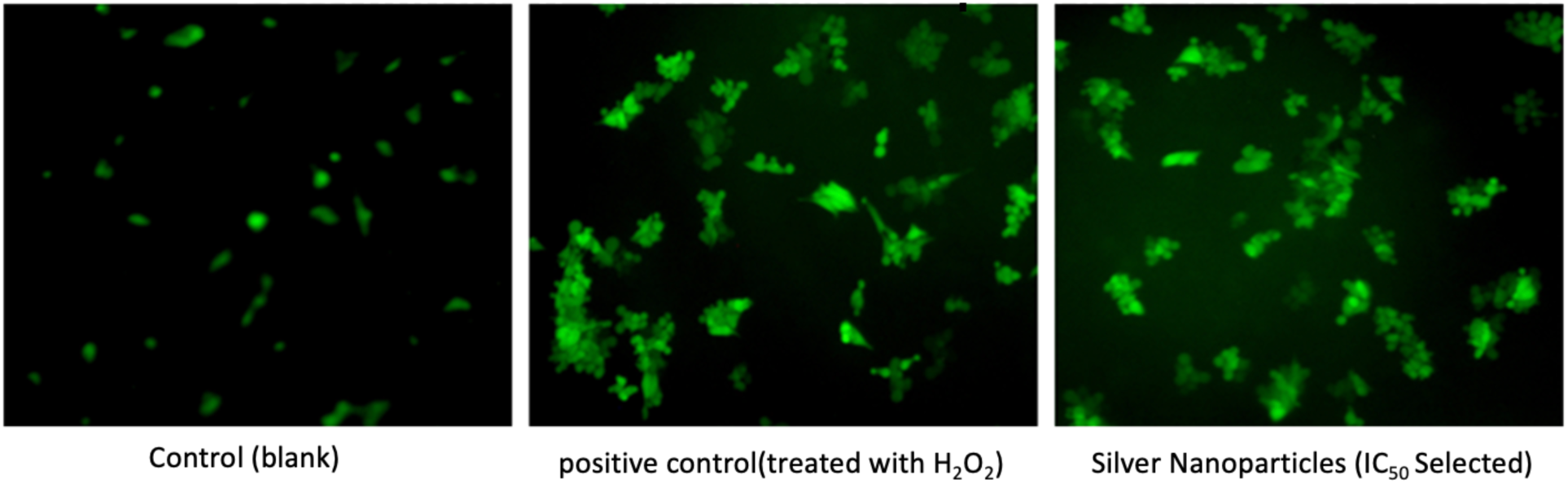
Reactive Oxygen Species production determination by 2՛,7՛-dichlorodihydrofluorescein diacetate: -SKOV3 cells fluorescent 10X magnification images. The control (Blank) has cell growth media and positive control has (0.03%) hydrogen peroxide within media. The cells were treated with of silver nanoparticles of the mentioned IC_50_ previously.

The reactive oxidizing molecules derived from ROS, have an impact on cell survival as they are endogenously produced (**Li et al., 2015**) They are defined as reactive molecules or ions that are formed by the incomplete one-electron reduction of oxygen. Superoxide (O^2-^), hydrogen peroxide (H_2_O_2_) and hydroxyl radicals (OH^-^) include the main types of ROS produced in the cell. The maximum concentration of ROS in the cells is considered to be produced by mitochondria (**Collins et al., 2012**).

#### 2.1.5 Molecular Docking

##### Binding site prediction

The active binding site of PARP1 protein (PDB ID 5DS3) was predicted by using the CASTp server. Thirty-one active pockets were identified, with the largest pocket having an area of 317.23 A° and volume of 385.35 A°. for further analysis, we have utilized UCSF chimera tool to eliminate cavities with zero openings resulting in the identification of only twelve cavities out of thirty-one active pockets, the amino acid residues involved in the active pocket are highlighted in **Figure 11**.

**Figure 11:**
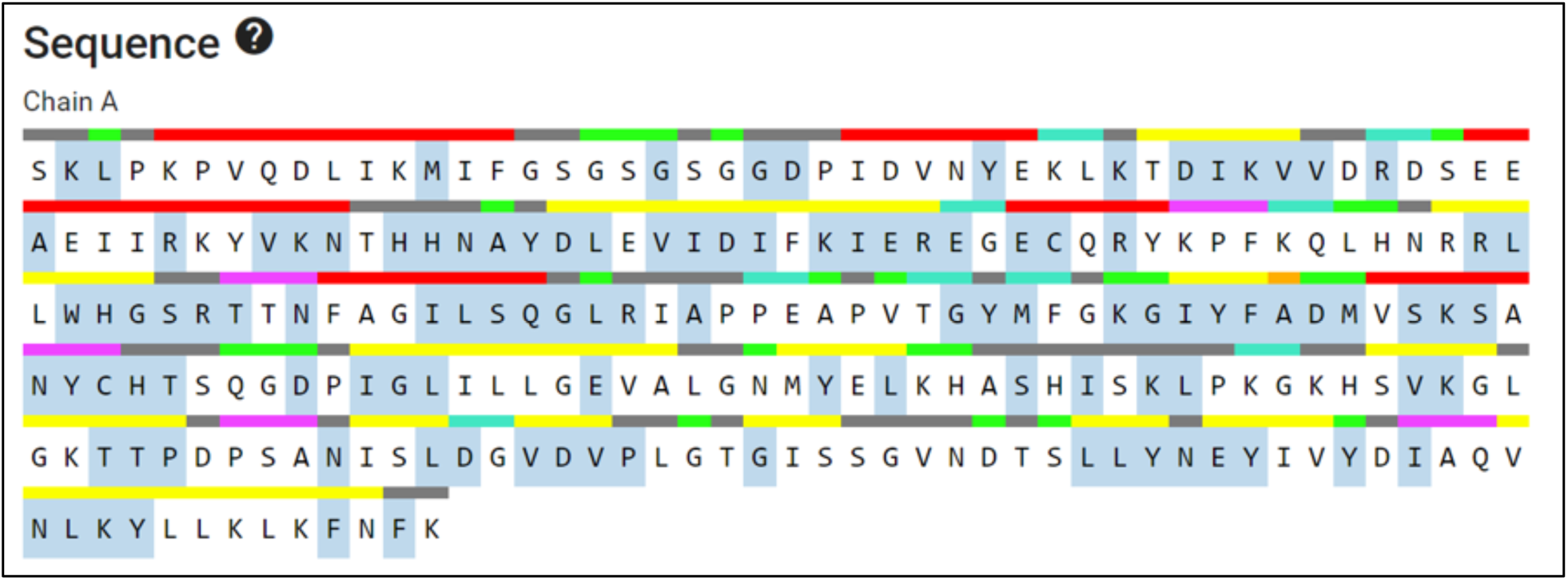
Active site prediction by CASTp

##### Interaction analysis

To elucidate the interaction between silver atoms and the targeted protein PARP1, molecular docking was performed using Autodock 4.2 and Cygwin terminal. In the docking process, the silver atom exhibited a binding energy of < 0 kcal/mol, indicating favorable free energy for interaction with the binding residues of the protein. We have also used optimized structures of Ag, Ag^+^ and Ag_3_ for molecular docking with PARP1 protein. The interaction analysis showed that Ag interacts with three amino acids of the PARP1 protein (**Figure 12**). For ionic silver, we have found that Ag^+^ interacted with the GLU844, ARG847, ALA995 and VAL997 amino acids of the PARP1 protein. Meanwhile, Ag_3_ shows the interaction with one active site residue ASN987 (**Table 2**). We have also analyzed the role of different phytochemicals present in the *S. bryopteris* that are responsible for reducing, capping and stabilization of AgNPs. The goal of molecular docking is to use a search algorithm approach to determine the binding mode of ligand to a protein. To achieve an optimized arrangement of both interacting molecules, with the aim of reducing the overall free energy of the system. The final predicted binding free energy (ΔG_bind_) is expressed through various factors such as dispersion and repulsion (ΔG_vdw_), hydrogen bonding (ΔG_hbond_), desolvation (ΔG_desolv_), electrostatic interactions (ΔG_elec_), torsional free energy (ΔG_tor_), the ultimate total internal energy (ΔG_total_), and the energy of the unbound system (ΔG_unb_). Therefore, gaining a comprehensive understanding of the fundamental principles governing the predicted binding free energy (ΔG_bind_) offers valuable insights into the nature of the various interactions that facilitate the docking of molecules (**Agarwal et al., 2016**). Based on the idea of the Lamarckian Genetic Algorithm (LGA), AutoDock 4.2 facilitates the analysis of protein-ligand interactions by identifying the lowest binding energy score indicate that the ligand has formed several interactions with the enzyme. The docking complex was retrieved in pdbqt file format from AutoDock Tools to visualize the interactions between proteins and ligands. We analysed the interaction between various phytochemicals and Poly (ADP-ribose) polymerase (**Table 3**). From the docking results, we have analysed various hit flavonoid compounds that were able to target the active site residues of PARP protein with promising ADME profiles. The chemical structures of these hit or lead compounds are shown in **Figure 13**. Interaction calculation between the ligands (Rhamnetin, Rutin) with the chosen protein (PARP1) was done using binding energy (i.e., docking score) (B.E.) kcal/mol) (**Figure 14**) and inhibition constant (nM), with interacting protein residues

**Figure 12:**
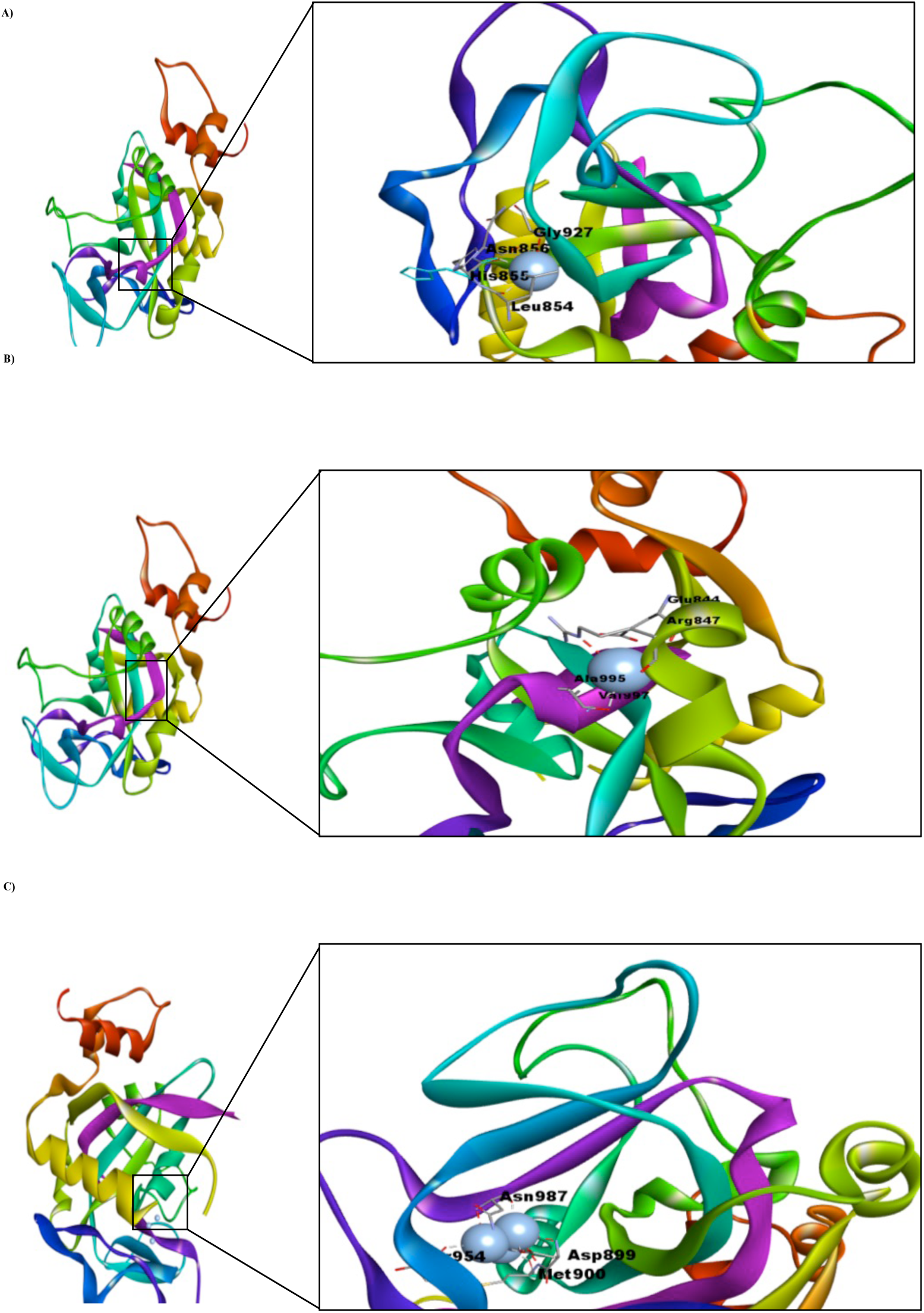
Molecular docking interactions of silver atoms (a) Ag, (b) Ag^+^, and (c) Ag_3_ with the amino residues of PARP1. The binding of the silver to the protein is illustrated on the left side, while the interactions between the amino acid residues and the silver atoms are depicted on the right side.

**Figure 13:**
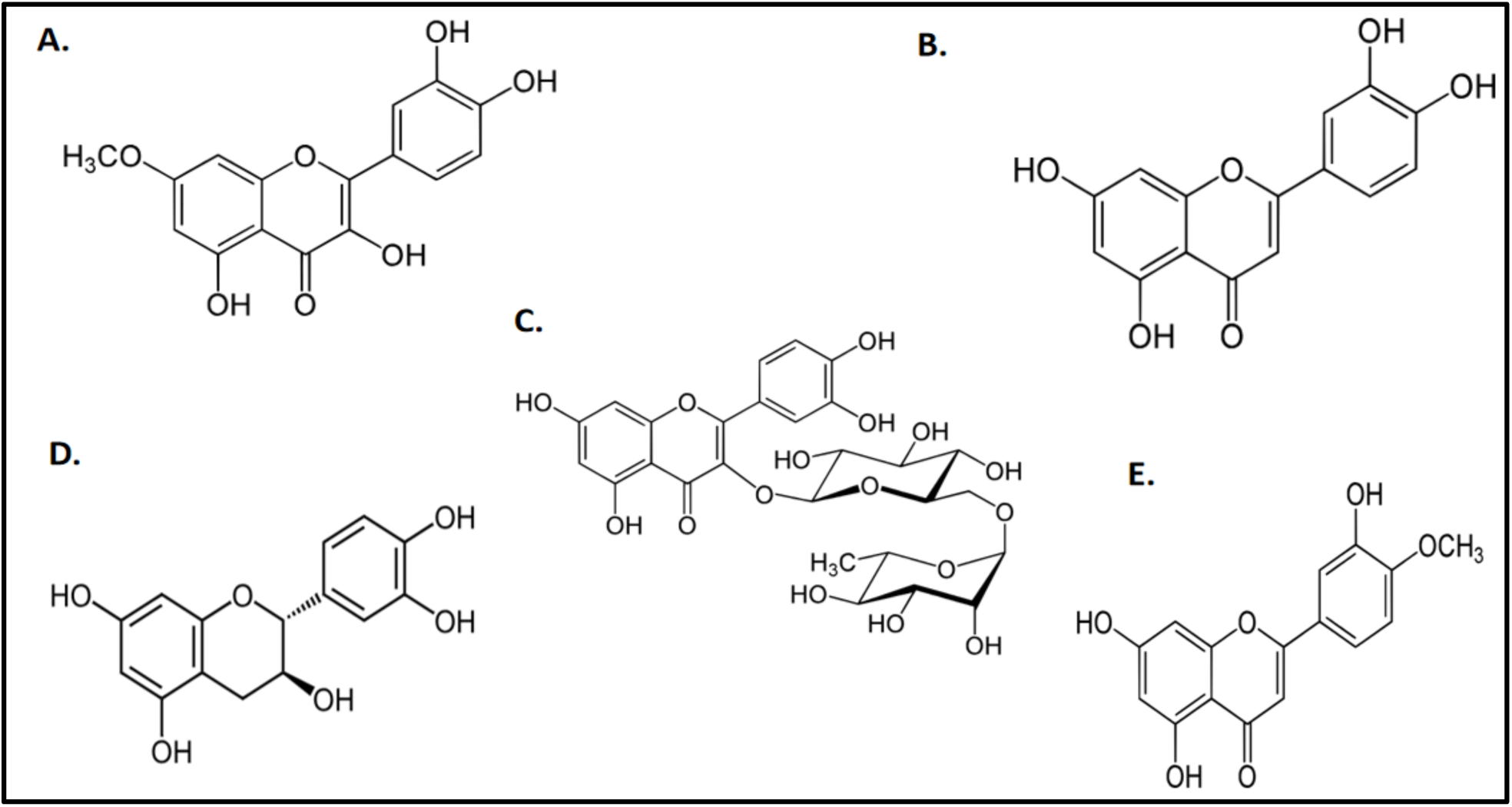
Structures of best hit flavonoids against PARP1 with favourable ADME properties (A) Rhamnetin (B) Luteolin (C) Rutin (D) Catechin (E) Diosmetin

**Figure 14:**
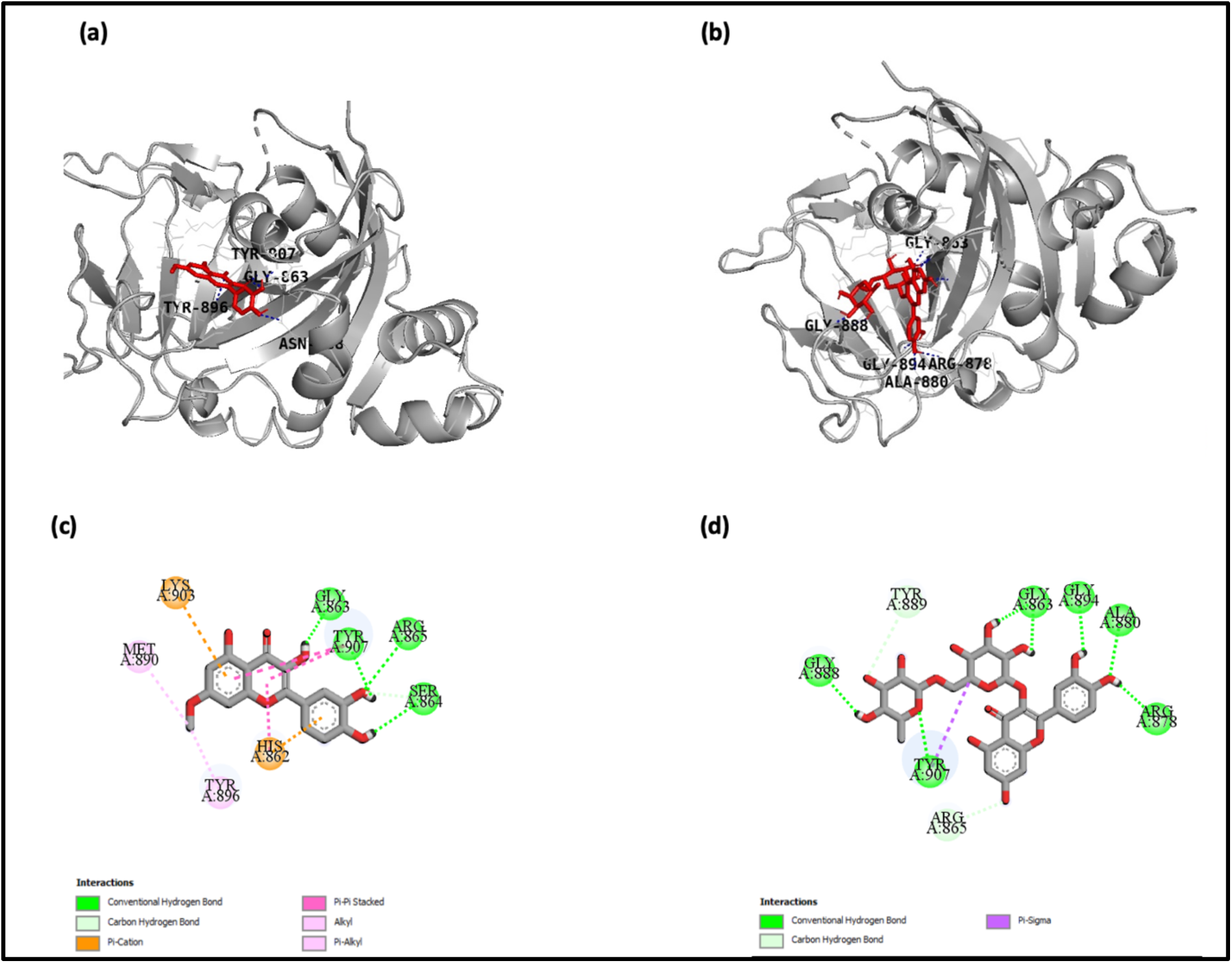
3D interaction established by (A) Rhamnetin, (B) Rutin; against PARP1, 2D interaction of (C) Rhamnetin (D) Rutin with PARP1.

**Table 2.** Interactions of Ag, Ag^+^ and Ag_3_ Cluster with amino acid residues of the binding site pocket of PARP1 protein.

| Macromolecule | Ligands | Binding energy (Kcal/mol) | Inhibition constant (mM) | Interacting Amino residues |
| --- | --- | --- | --- | --- |
| PARP1 | Ag | -0.21 | 700.74 mM | HIS855, LEU854 and GLY927 |
|  | Ag<sup>+</sup> | -0.18 | 736.01 mM | GLU844, ARG847, ALA955 and VAL997 |
|  | Ag<sub>3</sub> | -1.08 | 160.24 mM | ASP899, MET900, THR954 and ASN987 |

**Table 3.** Binding score of selected phytochemicals present in aqueous extract of *Selaginella bryopteris* against PARP1.

| Class | Name of Compound | Binding Energy (kcal/mol) | Inhibition Constant | Reference |
| --- | --- | --- | --- | --- |
| **Reference Ligand** | Olaparib | -15.15 | 7.4pM | (Cortez et al., 2018) |
| **Alkaloids** | 8-Hydroxyquinoline | -7.68 | 2.36 uM | (Adnan et al., 2021) |
|  | Guvacoline | -7.13 | 5.96 uM |  |
| **Flavonoids** | Rutin | -16.13 | 1.49 pM | (Adnan et al., 2021) |
|  | **Rhamnetin** | **-11.56** | **3.34 nM** |  |
|  | Catechin | -11.40 | 4.42 nM |  |
|  | Luteolin | -11.19 | 6.29 nM |  |
|  | Diosmetin | -10.83 | 11.58 nM |  |
|  | Kaempferol | -10.61 | 16.65 nM |  |
|  | Myo-inositol | -9.00 | 253.36 nM |  |
| **Biflavonoid** | 2,3-Dihydroamentoflavone | -17.31 | 205.83 fM | (Kunert et al., 2008) |
|  | Tetrahydroamentoflavone | -16.78 | 503.66 fM |  |
|  | Bilobetin | -16.73 | 545.08 fM |  |
|  | Amentoflavone | -16.67 | 608.68 fM |  |
|  | Sequoiaflavone | -15.93 | 2.10 pM |  |
|  | Heveaflavone | -15.81 | 2.57 pM |  |
|  | Sciadopitysin | -15.73 | 2.94 pM |  |
|  | Lanaroflavone | -14.79 | 14.35 pM |  |
|  | 2,3-Dihydrohinokiflavone | -13.77 | 80.65 pM |  |
|  | 2",3"-dihydroamentoflavone | -13.52 | 122.62 pM |  |

The binding energies of the phytocompounds ranged from -17.31 to -7.13 kcal/mol. The active site residues in PARP-1 are Trp 861, His862, Gly 863, Ser 864, Arg 865, Asn 868, Gly 876, Leu877, Arg 878, Phe 891, Ile 895, Tyr 896, Phe 897, Ala898, Lys 903, Ser 904, Tyr 907, Asn 987, Glu 988, Tyr989 (**Shridhar et al., 2021**). Phytocompound 2,3-Dihydroamentoflavone has the lowest energy -17.31 kcal/mol but shows unfavorable binding with a PARP-1 protein receptor. Rutin shows favorable binding interactions with this receptor, exhibiting a binding energy of -16.13 kcal/mol (**Table 3**). Notably, it forms a Pi-sigma bond with Tyr907 and establishes a hydrogen bond with Gly 863 from the selected active binding site (Figure 15 b, d). Subsequently, rhamnetin shows a docking score of -11.56 kcal/mol (**Table 3**) and shows interaction with His862, Gly 863, Ser 864, Arg 865, Tyr 896, Lys 903 and Tyr 907 from the selected active binding site (Figure 15 a, c). His 862 residues of PARP-1 proteins have ionization properties due to the presence of imidazole rings at active site. Analysis of PARP-1 protein through homology modeling has revealed two important intermolecular interactions and these interactions are found to be consistent with the results obtained from homology modelling of other superfamily members including Diphtheria toxin or endotoxin A (**Nilov et al., 2020**). Rhamnetin, a flavonoid phytocompound present in the extract of *S. bryopteris* can target the binding site of ADP and NAD^+^ at catalytic domain and shows interaction with the important reported residues of PARP protein including Gly 863, Tyr 907, Met890 Lys 903 (**Figure 14A and C**). Moreover, Rhamnetin has exhibits more promising ADME profiles as compared to rutin (**Table 4**). Additional investigation also reveals that the phytocompounds Catechin, Luteolin, and Diosmetin are also drug like candidates for ovarian cancer. Based on these results, we have identified Rhamnetin as the hit ligand molecule capable of targeting active site residues (Table 4).

**Figure 15:**
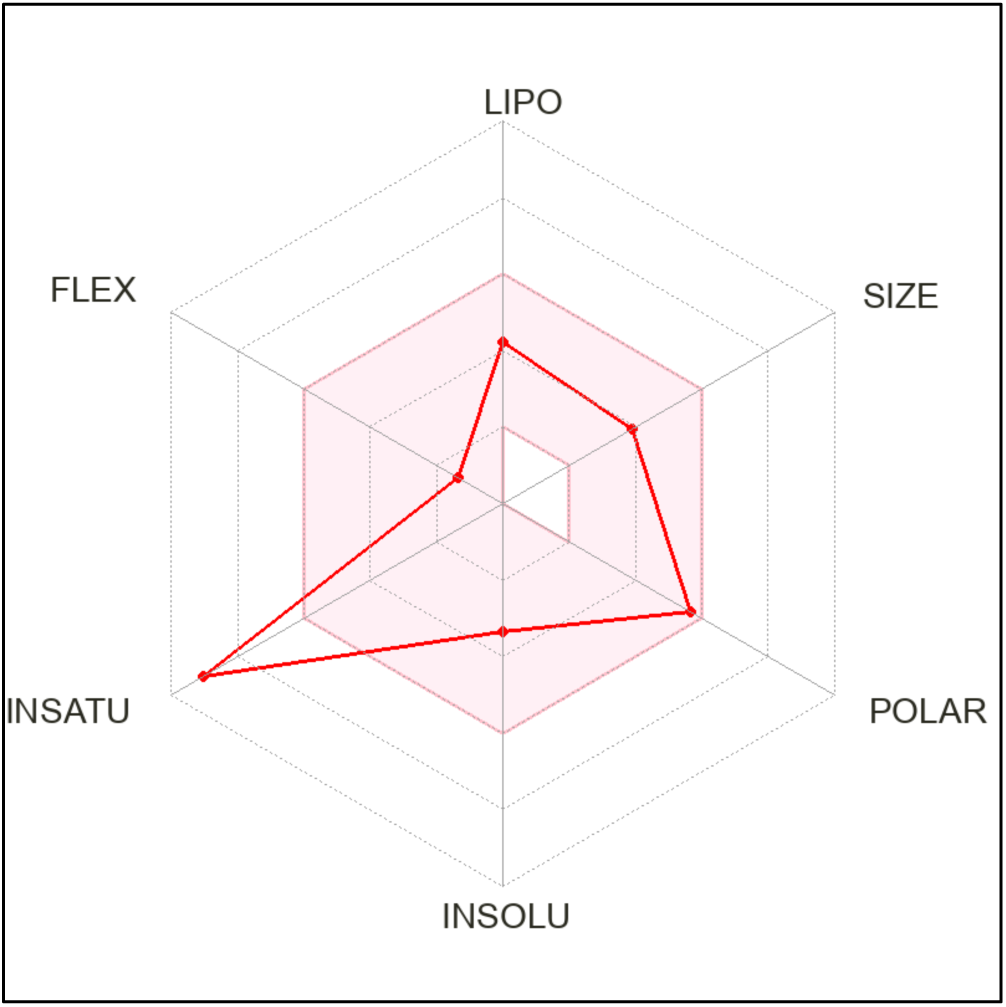
Bioavailability radar plot of Rhamnetin; LIPO (lipophilicity); XLOGP3 (-0.33), SIZE (molecular weight 316.26g/mol), POLAR (polarity 120.36 A°), INSOLU (insolubility -3.36), and FLEX (flexibility, number of rotatable bonds 6)

**Table 4.**
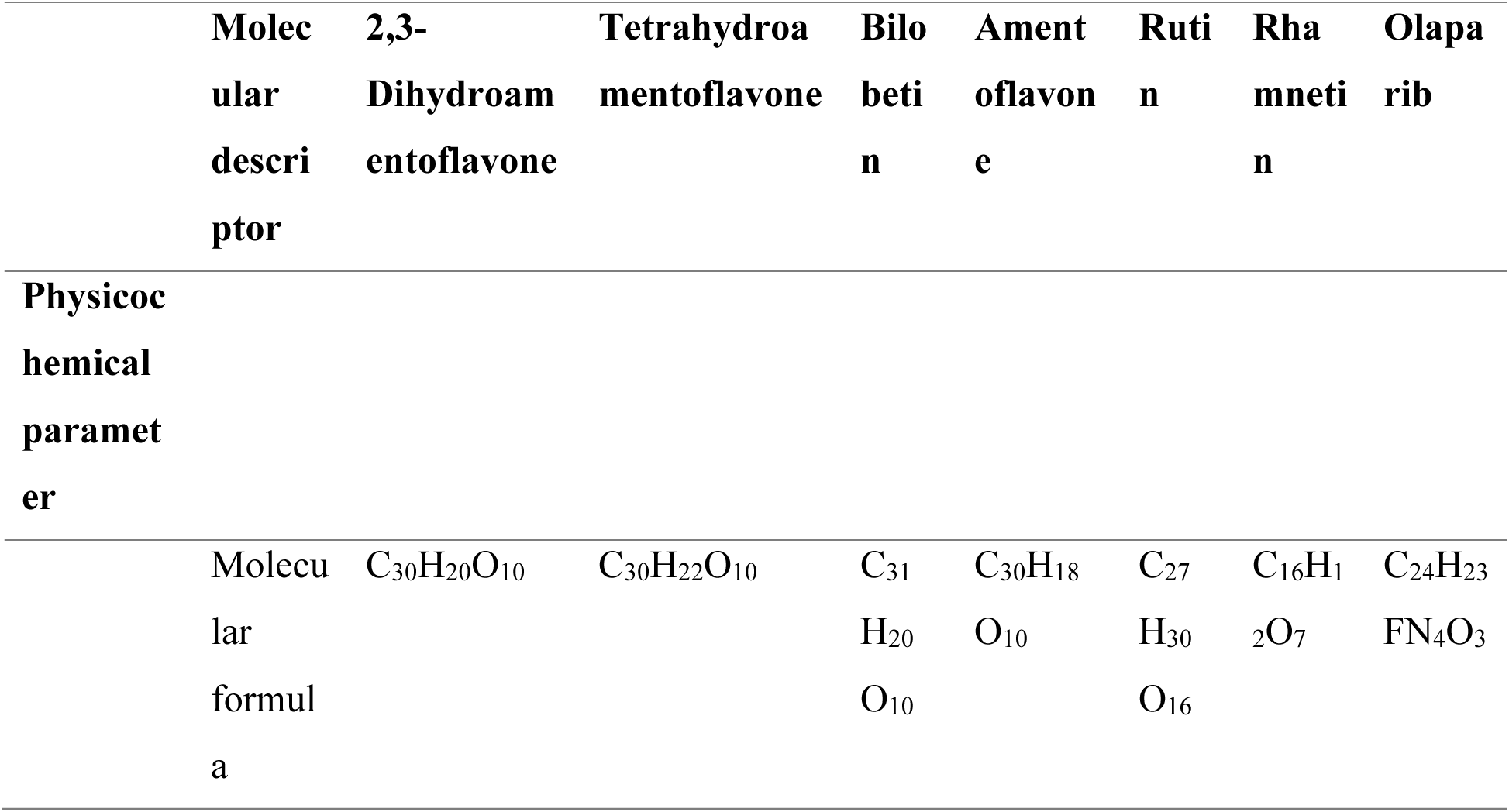
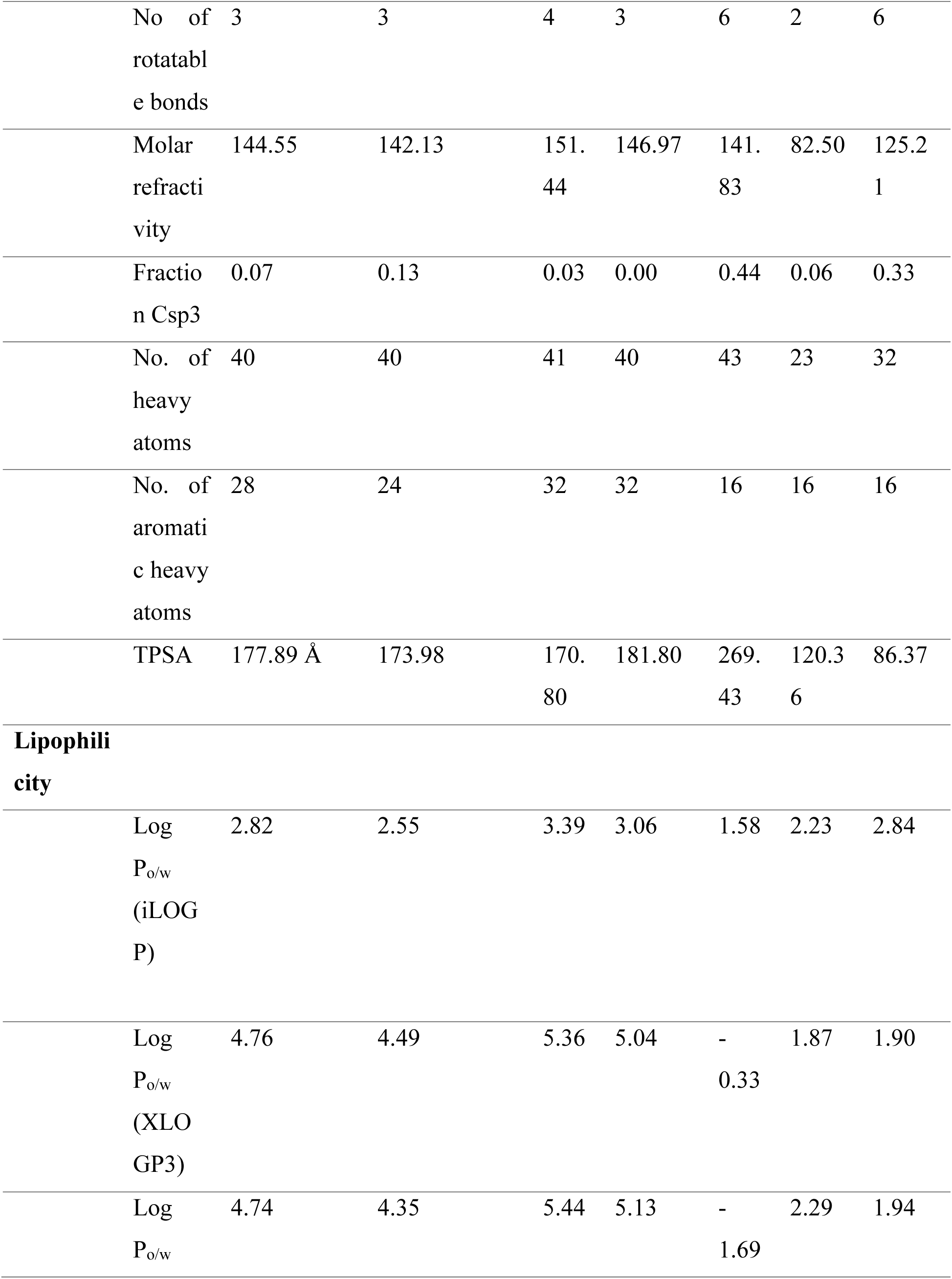
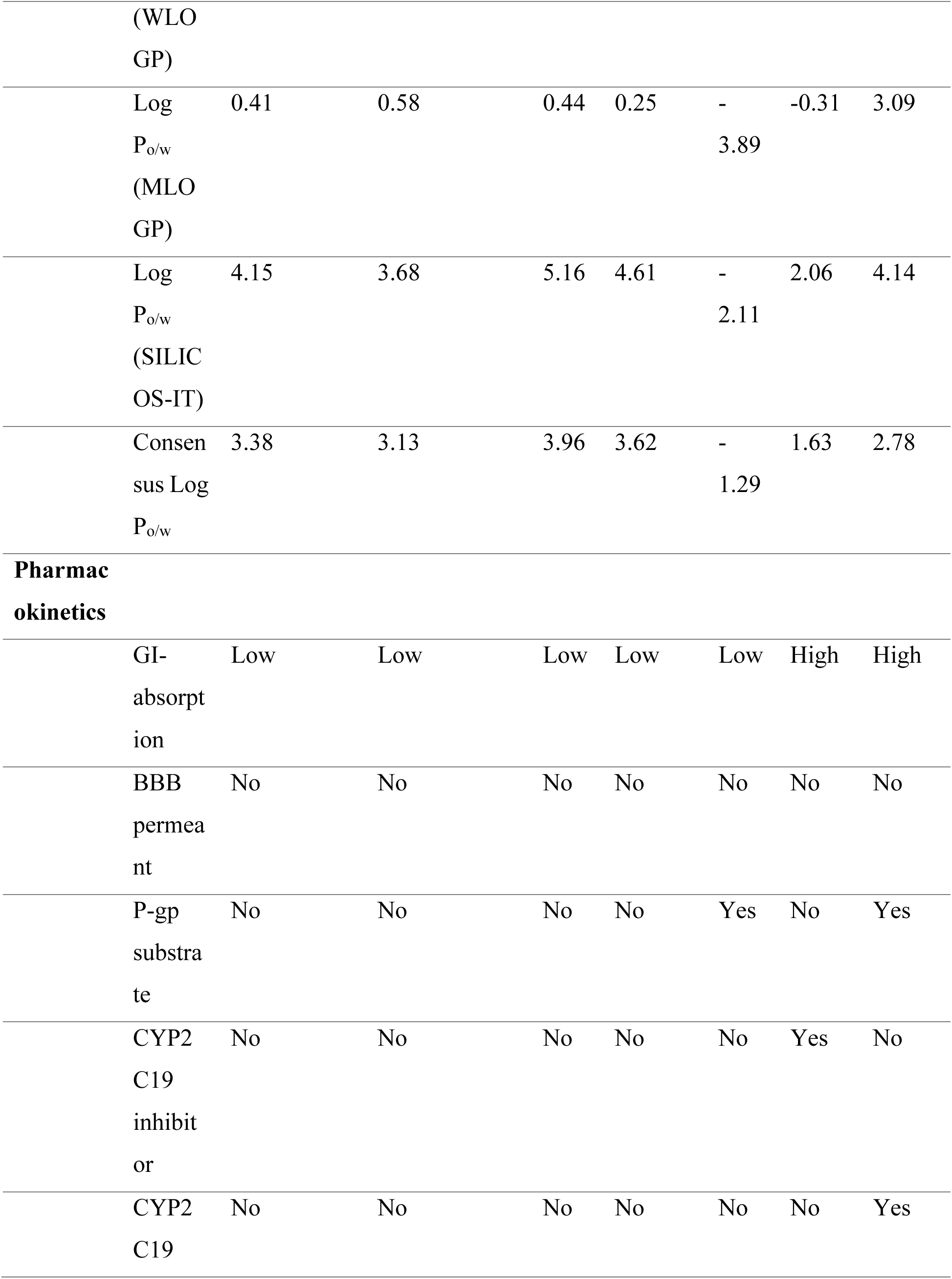
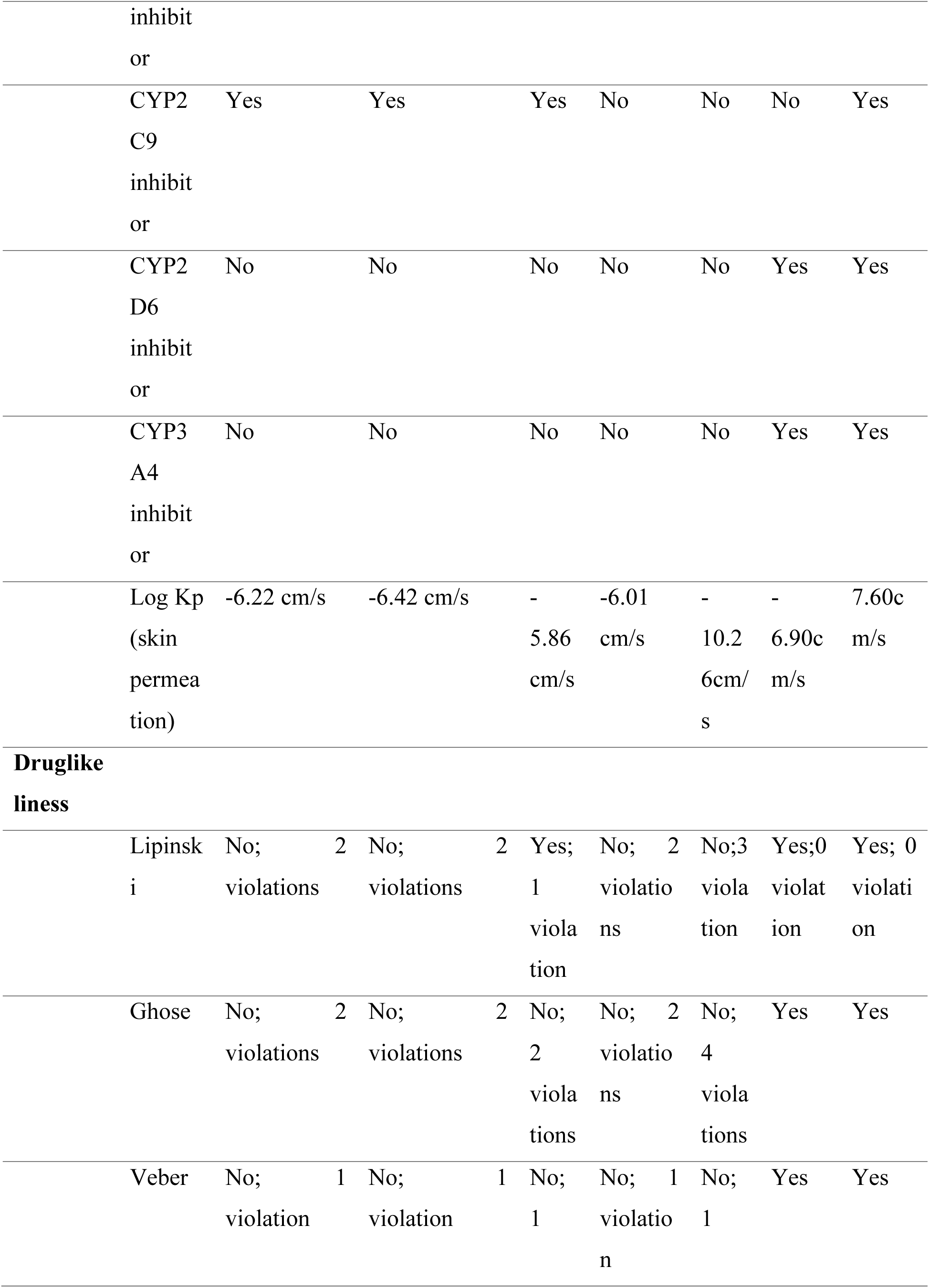
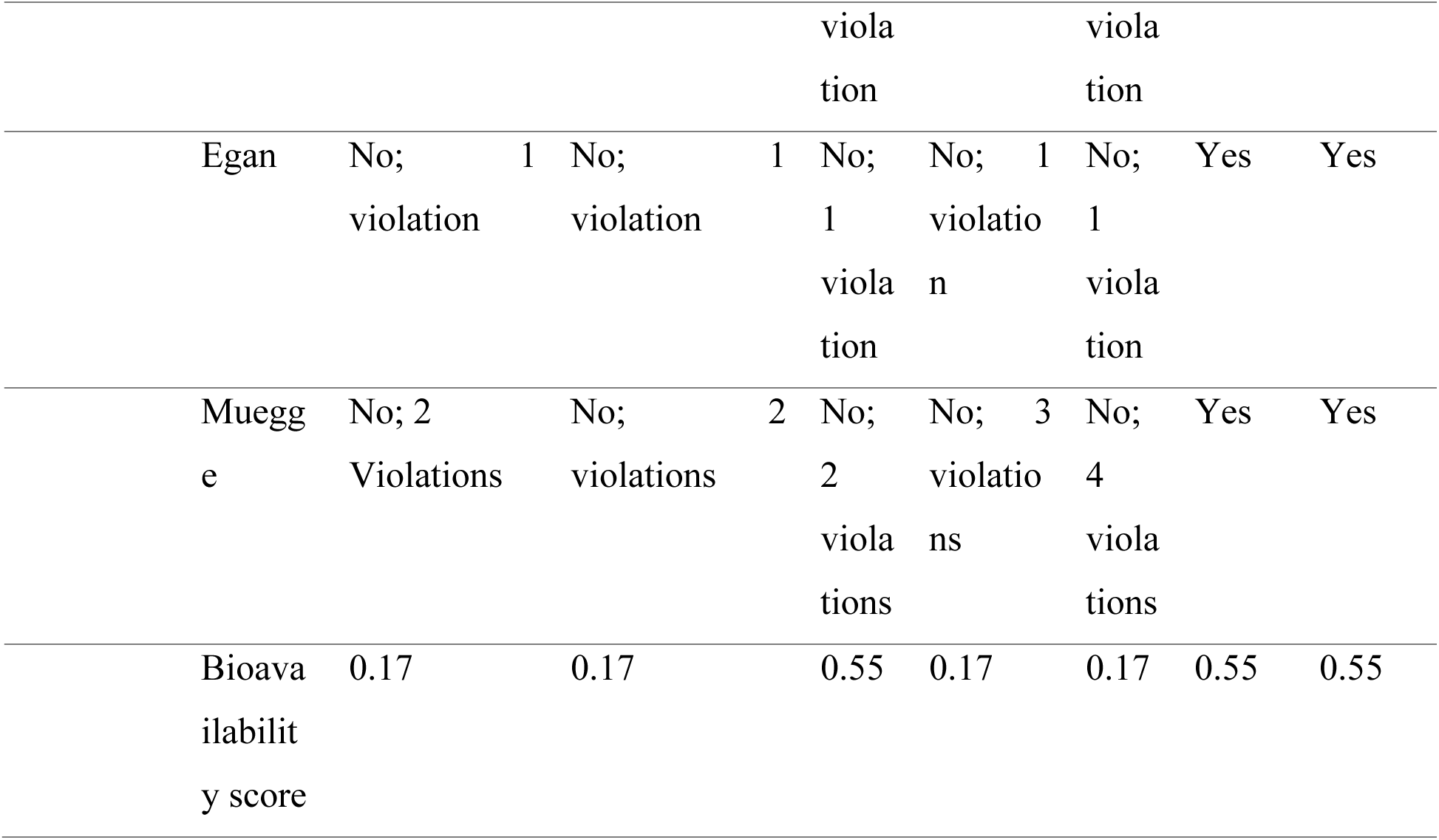
The ADME profile and drug-likeness of the top-ranked phytocompounds.

#### 2.1.6 Physicochemical and Pharmacokinetic Properties

The drug-likeness of the phytocompounds was evaluated by considering various ADME-related characteristics, which encompass pharmacokinetics, physiological parameters, water solubility, lipophilicity, and medicinal chemistry. The data in (**Table 4**) indicates that Rhamnetin exhibits the most promising potential as a drug candidate among the various compounds due to least violation. The prediction of ADME was obtained from Swiss ADME software, and their results are shown in **Table 4**. The number of rotatable bonds for the rhamnetin were less than 10 with hydrogen bond acceptor (<10) and hydrogen bond donor (<5). The molecular weight of rhamnetin was observed to be less than 500 KDa. Rhamnetin did not violate any Lipinski’s rule of five. The drug likeness of Rhamnetin is shown by the bioavailability radar plot in **Figure 15**. Light Pink region denotes the ideal range for each of the following characteristics: solubility: log S not greater than 6, saturation: fraction of carbons in the sp^3^ hybridization not less than 0.25, flexibility: no more than 9 rotatable bonds, polarity: TPSA between 20 and 130 A°, size: MW between 150 and 500g/ml, and lipophilicity: XlogP3 between -0.7 and +5.0.6) (Figure 15).

## 3. Materials and Methods

### 3.1 Materials

Dried *Selaginella bryopteris* was purchased from market, while the human ovarian cancerous cell line SKOV3 was obtained from the National Centre for Cell Science (NCCS), Pune, INDIA. AgNO_3_ was purchased from from Sigma Aldrich, which is located in St. Louis, MO, USA.

### 3.2 Methodology

#### 3.2.1 Collection and processing plant extract

*S. bryopteris* was washed thoroughly first with tap water followed by distilled water and was left to air dry for the next two days. Its leaves were then crushed to acquire a fine powder. The obtained leaf extract was carefully covered with aluminum foil and kept in cold, at 4°C. After mixing 0.25 grams of this leaf powder with 25 mL of distilled water, the mixture was stirred for an hour at room temperature using a magnetic stirrer. The process of obtaining aqueous leaf extract involved filtering using Whatman filter paper followed by centrifuging the filtrate for 10 minutes at 4700 rpm to eliminate any remaining plant residues. This leaf extract was used in the synthesis of AgNPs as a stabilizing and reducing agent.

#### 3.2.2 Silver nanoparticle synthesis using plant extract

After dissolving 0.0033g of AgNO_3_ in 10ml of distilled water, 10 ml of a 0.001M aqueous solution of silver nitrate was prepared and kept in an amber bottle to avoid the auto oxidation of Ag. 1mL of extracts of the *S. bryopteris* were taken and added to 9 mL of prepared 1mM solution of silver nitrate. The nanoparticle solution was maintained at room temperature for 24 hours at pH 7 to analyze by UV-Visible spectrophotometer (**Yadav et al., 2020**). After 48 hours, the silver nanoparticle suspensions were centrifuged at 12,000 rpm for 15 minutes and washed with distilled water at least three times to remove any undesired plant extract or waste. The change in color from transparent to brown indicated the formation of (AgNPs).

#### 3.2.3 Characterization of prepared silver nanoparticles (AgNPs)

In our previous study, we characterized biosynthesized silver nanoparticles using SEM and XRD, UV spectroscopy, TEM, and FTIR-ATR (**Yadav et al., 2020**). To ascertain the wavelengths of biosynthesized silver nanoparticles, an aliquot (2 mL) was extracted after 24 hours and was subjected to UV-spectroscopy Model UV-1800, Shimadzu, Japan, at a wavelength range of 200-800 nm. The observation of the distinct surface plasmon resonance (SPR) band corresponding to the biosynthesized AgNP validates the process of AgNP production. Further shape and size of AgNPs were determined by SEM equipped with EDX and TEM. For this purpose, dried AgNPs were sonicated for enough time, and then subjected onto platinum grid, and finally subjected to SEM. TEM was performed to determine the exact size of silver nanoparticles. Sonicated samples were loaded on a carbon-coated copper grid and was allowed to dry at room temperature and then subjected to transmission electron microscopy. Fourier-Transform Infrared Spectroscopy-Attenuated Reflectance (FTIR-ATR) was conducted for identification of the functional groups present in *S. bryopteris* leaf extracts. A Perkin Elmer spectrophotometer operating in transmission mode within the range of 400 -4000 cm^-1^ at a resolution of 4cm^-1^ was used to produce the FTIR-ATR spectra. The chemical composition of leaf extract of *S. bryopteris* was determined by using inductively coupled plasma mass spectrometry (ICP-MS) analysis by microwave digestion. The dried powder leaf extract was placed in a dry, clean Teflon microwave digestion vessel to which 2 mL of concentrated nitric acid and 6mL of concentrated hydrochloric acid was added. The samples were then subjected to digestion using a scientific microwave at 165°C for 10 minutes with the microwave irradiation of 1000 W for 15 minutes. After cooling, the sample was analyzed by using instrument Agilent 8900 Triple Quadrupole (model number G3665A) for the ICP-MS analysis.

#### 3.2.4. Stability of green synthesized silver nanoparticles at different pH and temperature

The pH stability of green synthesized silver nanoparticles was investigated by changing the pH of silver nanoparticles. For this, pH levels of 3, 5, 7, 8 and 9 were used. The pH of the solution was maintained by using freshly prepared 10% HNO_3_ and 0.1 M NaOH (**Mohammad Tahir et al., 2020; Summer et al., 2023; Mumtaz et al., 2023**). The heat stability of AgNPs was checked at different temperatures such as 3°C, 55°C, 75°C and at room temp (stock AgNP). UV-Vis spectrophotometer was used to observe the absorption spectra pattern of the reaction mixture after 24 h of incubation. Before measurement of UV-Visible spectra, samples were thoroughly mixed and vortexed. Each experimental sample was prepared in triplicates and results were tabulated.

## 4. In Silico Studies

### 4.1 Preparation of ligand

A list of 19 *S. bryopteris* phytochemical compounds that are well known and documented in literature data were selected. The chemical structure of Ag-Commercial Item Description (CID) No 23954 and Ag^+^ CID No 104755 were downloaded from PubChem compound database. For a more detailed study of the interaction between protein and ligand, AgNPs were modelled as Ag_3_ clusters containing three atoms of silver using Avogadro software (**Hanwell et al., 2012**). To investigate the mode of action of these phytocompounds against PARP proteins involved in DNA-damage repair in human ovarian cancer cell line SKOV3, molecular docking investigation was conducted by using AutoDock tools version 4.2 (**Morris et al., 2009**).

### 4.2. Unveiling Protein Binding Sites: Using CASTp and UCSF Chimera

The crucial areas of the protein that are utilized for docking with external ligands are called as pocket or binding sites were predicted by using CASTp server (**Tian et al., 2018**). The CASTp (http://sts.bioe.uic.edu/castp/) server displays the overall number of pockets as well as the amino acid residues involved in each pocket. Additionally, the pockets were further arranged by area and volume. Using UCSF Chimera, we visualized the result of CASTp server.

#### 4.2.1. Preparation of protein

RCSB Protein Data Bank (https://www.rcsb.org/) was used to obtain the structure of the PARP-1 protein (PDB 1D 5DS3) in PDB format. Using the ‘Open Babel 2.4’ programme, the 3D structures of a few chosen phytochemicals that were retrieved from PubChem in a sdf (particular file format) were converted to pdbqt format. Using AutoDock 4.2 and via AutoDock Tools 1.5.7 docking was carried out (**Morris et al., 2009**). The protein’s PDB structure was accessed using AutoDock Tools. Water molecules, heteroatoms and bound ligands were then removed from the protein. Polar hydrogen and Kollman charges were then added to the protein molecule. For the study, the grid parameter was chosen to encompass the active site of the protein. In this study, the grid spacing was set to 0.375 A° (as default) and center grid box were set to a x=5.447, y=36.952 and z=10.174, respectively. The number of grid points along was x, y and z dimension were set 80 x 80 x 90. The dimensions of x, y, and z were fixed to 80 x80 x90 grid points. The number of GA runs after auto grid finished were thirty. The default values for all other parameters were applied.

### 4.3 Molecular docking

Molecular docking was done to understand how ligands interact with specific proteins (enzymes). It is a useful and efficient technique for finding potential lead compounds in drug discovery. Virtual screening, where small molecules are docked into known protein structures, is an important part of this process. Through the process of virtual screening within databases of small molecules, it becomes feasible to discover novel candidate inhibitors for a specific target. Auto Dock tools make this process faster and cheaper to find new drug candidates. Auto Dock 4.2 is a popular docking program, that uses a Lamarckian genetic algorithm to determine the interaction of ligands with the protein and their binding energy, which represents how ligands are bound to the target protein (**Morris et al., 2009; Wadhwa et al., 2023**). Each docking software utilizes distinct search algorithms, which are the techniques employed for anticipating potential configurations of the binary complex.

### 4.4 Analysis of the best flavonoids’ drug disposition as potential therapeutic possibilities

SwissADME online programme was used to predict the pharmacokinetics and toxicity of the identified phytocompounds from the *S. bryopteris* extract (**Daina et al., 2017**).

It is crucial to remember that a compound’s ability to interact with enzymes or proteins by itself does not ensure that it will function as a drug. Drug-likeness studies evaluate a molecule’s bioavailability to predict its propensity to function as an oral medication. Understanding the various properties is crucial in drug design, as the presence of undesirable ADMET properties can fail a drug during clinical trials. Analysis by ADMET provides accurate information regarding the drug-like characteristics of the phytochemical present in *S. bryopteris* including numerical values for parameters such as permeation of the blood-brain barrier, toxicity test and gastrointestinal absorption.

## 5. Anticancer activity of Silver Nanoparticles

### 5.1 Cell line

National Centre for Cell Service (NCCS), Pune, India provided the cancer cell line SKOV3. Roswell Park Memorial Institute Medium 1640, supplemented with 10% (v/v) foetal bovine serum and 1% (v/v) streptomycin, was used to cultivate the cells. The cells were incubated at 37 °C in CO2 (5%) atmosphere. The cultures were properly cleaned in phosphate-buffered saline (PBS, pH 7.4) and then detached using a 0.25% trypsin-ethylenediaminetetraacetic acid (EDTA) solution prior to each experiment.

### 5.2 MTT ASSAY (Invitro cytotoxicity analysis)

MTT test was used to assess cytotoxicity of silver nanoparticles on an ovarian cell line (**Denizot and Lang, 1986**). The cells at the exponential phase of development, corresponding to around 1X 10^5^ mL cells, were sown into 96 wells of a covered polystyrene plate. A 24-hour incubation period was conducted at at 37°C with CO_2_ (5%). The next day, three duplicates of various drug concentrations ranging from 200-1000µg/ml were put to the microtiter plate. The MTT reagent (5mg/ml) was added to wells and then incubated for an additional three hours after the first day of incubation. After three hours, the formazan crystals were allowed to dissolve in fifty microliters of isopropanol, and the measurement of absorbance was done using a Spectrostar-Nano plate reader ten minutes later at 620 nm. The control wells without any drug treatment were considered as control. The formula for calculating percentage cytotoxicity was (U-T)/U x 100, where U is the cells that were left untreated (control) and T is the mean optical density of the cells that were treated with various drug dosages.

### 5.3 Detection of intracellular Reactive Oxygen Species (ROS)

SKOV3 cells were grown on cover slips in a 35 mm petri plates in a CO_2_ incubator at 37°C for 24 hours. After 24 hours, cells were treated with IC_50_ values of silver nanoparticles for 24 hours. After incubation, cells were treated with H_2_O_2_ (0.03%) for 30 min (positive control) along with a negative control. The cells were treated twice with PBS and then incubated in medium containing 10 µM 2՛,7՛-dichlorofluorescein diacetate for 30 minutes at 37°C, 5% CO_2_ in the dark. Immediately after incubation, the cells were washed three times with serum-free medium to remove extracellular 2՛,7՛-dichlorofluorescein diacetate. The slides were prepared by inverting the coverslips having cells on cleaned slides on 20% glycerol drop and observed under 10X magnification by fluorescence microscope, Olympus, excitation at 480 nm and emission at 526 nm.

## Discussion

Utilizing plant derived compounds has been considered extensively for the treatment of cancer. Flavonoids, the secondary metabolites responsible for the color and flavours of vegetables, have already been acknowledged for their therapeutic potential against various health conditions. New, effective, and safe inhibitors are actively sought due to the limited effectiveness of current therapies, particularly those that target multiple receptor sites without causing undesirable effects. *Invitro* approaches made us conclude that the silver nanoparticles, which were biosynthesized utilizing the leaf extract derived from *S. bryopteris*, exhibit potent anti-cancer activity against the SKOV3 human ovarian cancer cell line. Further by *insilico* analysis we conclude that rhamnetin present in the aqueous leaf extract of *S. bryopteris* can be considered as an interesting scaffold for the development of the inhibitor of PARP protein to treat ovarian cancer in women. PARP inhibitors (PARPi) are promising therapeutic agents. Furthermore, we have also examined the formation of ROS by silver nanoparticles that enhances anticancer activity by reacting with cellular biomolecules. *S. bryopteris* is a model system for genetic studies and has an array of therapeutic properties. It’s conservation in changing natural habitats for ecological maintenance and human use is a primary step required for future research studies. In this context, examining phytochemicals derived from *S. bryopteris* like flavonoids is a logical approach. *Insilico* studies demonstrated that rhamnetin exhibits good safety profile and pharmacokinetics which makes it a potential drug candidate for ovarian cancer treatment. The use of niraparib, as PARPi has led to increased fatigue, anemia, thrombocytopenia, vomiting, nausea, decreased platelet count. The mitigation of these side effects of PARPi can be overcome by the use of phytochemicals which are generally made by plants for their defense mechanisms including terpenoids (lycopene, luteolin), polyphenols (flavonoids, catechins, quercetin, anthocyanins, curcumin, proanthocyanidins) and they can be used to treat ovarian cancer patients with BRCA mutations. Many phytochemicals including sulphur containing compounds (sulforaphane, allicin,) and above-mentioned phytochemicals are mostly derived from plants, fruits, vegetables and exhibit high safety profile. An in-depth understanding of the mechanism of phytochemicals targeting the ovarian cancer treatment is still unknown. In an equivalent way, further research using isolated rhamnetin is essential to thoroughly assess each of its characteristics. It is imperative to pursue studies for gathering additional data and enhance the utilization and versatility of rhamnetin like phytocompounds. Our future research will pave the way for developing new-generation therapeutic drugs or nanomedicines against various cancer types. With this, silver nanoparticles can be used in targeted drug delivery or as a carrier to transport chemical drugs and their controlled release. The modification of silver nanoparticles as drug carriers can help medical communities in the treatment of cancer. It can be anticipated that future clinical research will offer new insights into more effective targeted therapies.

## Abbreviations

PARPs: Poly ADP-ribose polymerases
PARPi: PARP inhibitor
AgNP: Silver nanoparticles
SKOV3: human ovarian cancer cell line
MTT: 3-[4,5-dimethylthiazol-2-yl]-2,5 diphenyl tetrazolium bromide
EOC: Epithelial ovarian cancer
MAPK: mitogen activated protein kinase
BRAF: v-Raf murine sarcoma viral oncogene homolog B
KRAS: Kirsten rat sarcoma
CDKN2A: Cyclin dependent kinase inhibitor 2A
PIK3CA: Phosphatidylinositol-4,5-bisphosphate 3- kinase catalytic alpha
TP53: Tumor protein 53
ARID1A: AT-rich interaction domain 1A
ERB2: v-erb-b2-avian erythroblastic leukemia viral oncogene homolog 2
HER2: Human epidermal growth factor receptor 2
NPs: Nanoparticles
MDR: Multi drug resistant
ARTD17: ADP- ribosyl transferase diphtheria-toxin-like proteins
BER: Base excision repair
SSBs: Single strand breaks
DSBs: Double strand breaks
HR: Homologous recombination
NHEJ: Non homologous end joining
BRCA1: Breast cancer gene 1
BRCA2: Breast cancer gene 2
UV-Vis: Ultraviolet-visible spectroscopy
SPR: Surface plasmon resonance
ADP: Adenosine diphosphate
DBD: DNA binding domain
AMD: Auto modification domain
CD: Catalytic domain
FTIR: Fourier Transform Infrared Spectroscopy
nm: nanometer
cm -1: Spectroscopic wavenumber
CH_3_: Methyl group
C-H: Carbon Hydrogen bond
C=N: Carbon Nitrogen bond
N-H: Nitrogen Hydrogen bond
O-H: Oxygen Hydrogen bond
μg/ml: microgram/millilitre
IC 50: Inhibitory concentration
LGA: Lamarckian Genetic Algorithm
PDBQT: Protein data bank partial charge atom type
Kcal/mol: Kilo calorie per mol
nM: Nanomolar
μM: Micromolar
pM: Picomolar
fM: Femtomolar
ADME: Absorption - Distribution - Metabolism and Excretion
BE: Binding energy
ROS: Reactive oxygen species
NCCS: National Centre for Cell Science
RT: Room temperature
rpm: Revolution per minute
FTIR-ATR: Fourier Transform Infrared Spectroscopy-Attenuated Total Reflectance
FBS: Fetal bovine serum
PBS: Phosphate buffer saline

## Author Contributions

Conceptualization, A.D., K.W., N.Kr., C.P., A.K., and N.K.; methodology, A.K., A.D., N.K., N.Kr., K.W., C.P; Botany, C.P., N.K; software, N.Kr.; validation, N.K., A.D., K.W., N.Kr., and A.K.; formal analysis, A.D., K.W., N.Kr., C.P., N.K.; investigation, draft preparation, K.W., N.Kr., C.P.; images, K.W., N.Kr., C.P., N.K; writing—review and editing, N.Kr., K.W., C.P., N.K; visualization, K.W., N.Kr., C.P., N.K; supervision, N.K.

## Ethics approval and consent to participate

No animals have been harmed and no ethics have been violated.

## Availability of data and materials

Not applicable.

## Financial Statement

The author KW wishes to acknowledge the University Grant Commission (UGC) for providing financial support in the form of Senior Research Fellowship (SRF).

## Graphical abstract

Schematic representation of anticancer activities of *S. bryopteris* derived silver nanoparticles against ovarian cancer associated PARP-1 protein.

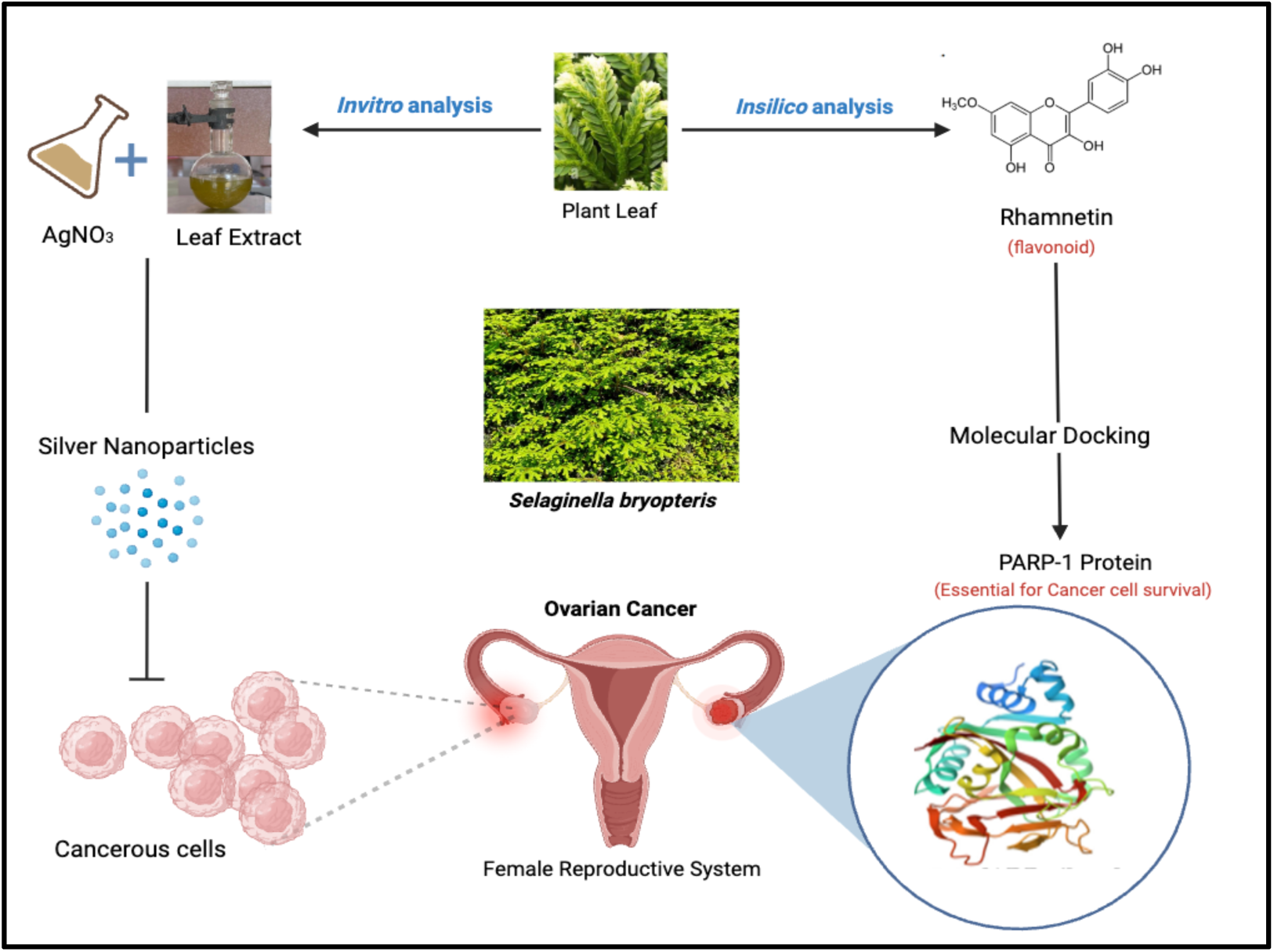

## References

Keyvani V, Kheradmand N, Navaei ZN, Mollazadeh S, Esmaeili SA. Epidemiological trends and risk factors of gynecological cancers: an update. Medical Oncology. 2023 Feb 9;40(3):93. 10.1007/s12032-023-01957-3

Shafabakhsh R, Asemi Z. Quercetin: a natural compound for ovarian cancer treatment. Journal of ovarian research. 2019 Dec; 12:1–9. 10.1186/s13048-019-0530-4

Berek JS, Kehoe ST, Kumar L, Friedlander M. Cancer of the ovary, fallopian tube, and peritoneum. International journal of gynecology & obstetrics. 2018 Oct; 143:59–78. 10.0002/ijgo/.12614

Kroeger Jr PT, Drapkin R. Pathogenesis and heterogeneity of ovarian cancer. Current opinion in obstetrics & gynecology. 2017 Feb;29(1):26. 10.1097/GCO.0000000000000340

Rojas V, Hirshfield KM, Ganesan S, Rodriguez-Rodriguez L. Molecular characterization of epithelial ovarian cancer: implications for diagnosis and treatment. International journal of molecular sciences. 2016 Dec 15;17(12):2113. 10.3390/ijms17122113.

Yeung TL, Leung CS, Yip KP, Au Yeung CL, Wong ST, Mok SC. Cellular and molecular processes in ovarian cancer metastasis. A review in the theme: cell and molecular processes in cancer metastasis. American Journal of Physiology-Cell Physiology. 2015 Oct 1;309(7):C444–56. 10.1152/ajpcell.00188.2015

Summer, M., Hussain, T., Ali, S., Khan, R. R. M., Muhammad, G., & Liaqat, I. (2024a). Exploring the underlying modes of organic nanoparticles in diagnosis, prevention, and treatment of cancer: a review from drug delivery to toxicity. International Journal of Polymeric Materials and Polymeric Biomaterials, 1–17.

Yin M, Xu X, Han H, Dai J, Sun R, Yang L, Xie J, Wang Y. Preparation of triangular silver nanoparticles and their biological effects in the treatment of ovarian cancer. Journal of Ovarian Research. 2022 Nov 21;15(1):121. 10.1186/s13048-022-01056-3

Pearce A, Haas M, Viney R, Pearson SA, Haywood P, Brown C, Ward R. Incidence and severity of self-reported chemotherapy side effects in routine care: A prospective cohort study. PloS one. 2017 Oct 10;12(10): e0184360. 10.1371/journal.pone.0184360

Torino F, Barnabei A, Paragliola R, Baldelli R, Appetecchia M, Corsello SM. Thyroid dysfunction as an unintended side effect of anticancer drugs. Thyroid. 2013 Nov 1;23(11):1345–66. 10.1089/thy.2013.0241

Anand U, Dey A, Chandel AK, Sanyal R, Mishra A, Pandey DK, De Falco V, Upadhyay A, Kandimalla R, Chaudhary A, Dhanjal JK. Cancer chemotherapy and beyond: Current status, drug candidates, associated risks and progress in targeted therapeutics. Genes & Diseases. 2022 Mar 18. 10.1016/j.gendis.2022.02.007

Ratan ZA, Haidere MF, Nurunnabi MD, Shahriar SM, Ahammad AS, Shim YY, Reaney MJ, Cho JY. Green chemistry synthesis of silver nanoparticles and their potential anticancer effects. Cancers. 2020 Apr 1;12(4):855. 10.3390/cancers12040855

Mohammad ZH, Ahmad F, Ibrahim SA, Zaidi S. Application of nanotechnology in different aspects of the food industry. Discover Food. 2022 Mar 22;2(1):12. 10.1007/s44187-022-00013-9

Summer, M., Ali, S., Tahir, H. M., Abaidullah, R., Fiaz, U., Mumtaz, S., … & Farooq, M. A. (2024b). Mode of action of biogenic silver, zinc, copper, titanium and cobalt nanoparticles against antibiotics resistant pathogens. Journal of Inorganic and Organometallic Polymers and Materials, 34(4), 1417–1451.

Summer, M., Ashraf, R., Ali, S., Bach, H., Noor, S., Noor, Q., … & Khan, R. R. M. (2024c). Inflammatory response of nanoparticles: mechanisms, consequences, and strategies for mitigation. Chemosphere, 142826.

Iravani S, Korbekandi H, Mirmohammadi SV, Zolfaghari B. Synthesis of silver nanoparticles: chemical, physical and biological methods. Research in pharmaceutical sciences. 2014 Nov;9(6):385. PMID: 26339255, PMCID: PMC4326978

Mandal D, Bolander ME, Mukhopadhyay D, Sarkar G, Mukherjee P. The use of microorganisms for the formation of metal nanoparticles and their application. Applied microbiology and biotechnology. 2006 Jan; 69:485–92. 10.1007/s00253-005-0179-3

Gericke M, Pinches A. Biological synthesis of metal nanoparticles. Hydrometallurgy. 2006 Sep 1;83(1-4):132–40. 10.1016/j.hydromet.2006.03.019

Zulfiqar, Z., Khan, R. R. M., Summer, M., Saeed, Z., Pervaiz, M., Rasheed, S., … & Ishaq, S. (2024). Plant-mediated green synthesis of silver nanoparticles: synthesis, characterization, biological applications, and toxicological considerations: a review. Biocatalysis and Agricultural Biotechnology, 103121.

Abbas, Z., Irshad, M., Ali, S., Summer, M., Rasheed, A., & Jawad, M. (2024). Radical scavenging potential of spectrophotometric, spectroscopic, microscopic, and EDX observed zinc oxide nanoparticles from leaves, buds, and flowers extract of Bauhinia Variegata Linn: A thorough comparative insight. Microscopy Research and Technique.

Mishra S, Singh HB. Biosynthesized silver nanoparticles as a nanoweapon against phytopathogens: exploring their scope and potential in agriculture. Applied microbiology and biotechnology. 2015 Feb; 99:1097–107. 10.1007/s00253-014-6296-0

Choi YJ, Park JH, Han JW, Kim E, Jae-Wook O, Lee SY, Kim JH, Gurunathan S. Differential cytotoxic potential of silver nanoparticles in human ovarian cancer cells and ovarian cancer stem cells. International journal of molecular sciences. 2016 Dec 12;17(12):2077. 10.3390/ijms17122077

Wadhwa K, Kaur H, Kapoor N, Ghorai SM, Gupta R, Sahgal A. A systematic review on antimicrobial activities of green synthesized Selaginella silver nanoparticles. Expert Reviews in Molecular Medicine. 2023a Aug 3:1–4. 10.1017/erm.2023.21

Rahimi, F., Shahraki, S., Hajinezhad, M. R., Fathi-Karkan, S., Mirinejad, S., Sargazi, S., … & Saravani, R. (2024). Synthesis, characterization, and toxicity assessments of Silymarin-loaded Ni-Fe Metal-organic frameworks: Evidence from in vitro and in vivo evaluations. Journal of Drug Delivery Science and Technology, 92, 105372. 10.1016/j.jddst.2024.105372

Huang, S., Yuan, J., Xie, Y., Qing, K., Shi, Z., Chen, G., … & Zhou, W. (2023). Targeting nano-regulator based on metal–organic frameworks for enhanced immunotherapy of bone metastatic prostate cancer. Cancer Nanotechnology, 14(1), 43. 10.1186/s12645-023-00200-y

Bai, R., Zhu, J., Bai, Z., Mao, Q., Zhang, Y., Hui, Z., … & Xie, T. (2022). Second generation β-elemene nitric oxide derivatives with reasonable linkers: potential hybrids against malignant brain glioma. Journal of Enzyme Inhibition and Medicinal Chemistry, 37(1), 379–385. 10.1080/14756366.2021.2016734

Wei, S., Sun, T., Du, J., Zhang, B., Xiang, D., & Li, W. (2018). Xanthohumol, a prenylated flavonoid from Hops, exerts anticancer effects against gastric cancer in vitro. Oncology reports, 40(6), 3213–3222. 10.3892/or.2018.6723

Yadav V, Kapoor N, Ghorai SM. Green Synthesis of Silver Nanoparticles from Aqueous Leaf Extract of. Current Bioactive Compounds. 2020 Jun 1;16(4):449–59. 10.2174/1573407215666181122121039

Pandey V, Ranjan S, Deeba F, Pandey AK, Singh R, Shirke PA, Pathre UV. Desiccation-induced physiological and biochemical changes in resurrection plant, Selaginella bryopteris. Journal of Plant Physiology. 2010 Nov 1;167(16):1351–9. 10.1016/j.jplph.2010.05.001

Antony R, Thomas R. A mini review on medicinal properties of the resurrecting plant Selaginella bryopteris (Sanjeevani). International Journal of Pharmacy & Life Sciences. 2011 Jul 1;2(7). ISSN: 0976-7126

Sah NK, Singh SN, Sahdev S, Banerji S, Jha V, Khan Z, Hasnain SE. Indian herb ‘Sanjeevani’ (Selaginella bryopteris) can promote growth and protect against heat shock and apoptotic activities of ultra violet and oxidative stress. Journal of Biosciences. 2005 Sep; 30:499–505. 10.1007/BF02703724

Schulz C, Little DP, Stevenson DW, Bauer D, Moloney C, Stützel T. An overview of the morphology, anatomy, and life cycle of a new model species: the lycophyte Selaginella apoda (L.) Spring. International Journal of Plant Sciences. 2010 Sep;171(7):693–712. 10.1086/654902

Paswan SK, Srivastava S, Rao CV. Wound healing activity of ethanolic extract of Selaginella bryopteris on rats. Pharmacognosy Journal. 2020;12(2). 10.5530/pj.2020.12.53

Gautam, A., Kumar, V., Azmi, L., Rao, C. V., Khan, M. M., Mukhtar, B., … & Alam, A. (2023). Wound healing activity of the flavonoid-enriched fraction of Selaginella bryopteris Linn. against streptozocin-induced diabetes in rats. Separations, 10(3), 166.

Lu AZ, Abo R, Ren Y, Gui B, Mo JR, Blackwell D, Wigle T, Keilhack H, Niepel M. Enabling drug discovery for the PARP protein family through the detection of mono-ADP-ribosylation. Biochemical Pharmacology. 2019 Sep 1; 167:97–106. 10.1016/j.bcp.2019.05.007

Bürkle A, Virág L. Poly (adp-ribose): Paradigms and paradoxes. Molecular aspects of medicine. 2013 Dec 1;34(6):1046–65. 10.1016/j.mam.2012.12.010

Bürkle A. Poly (ADP-ribose) The most elaborate metabolite of NAD+. The FEBS journal. 2005 Sep;272(18):4576–89. 10.1111/j.1742-4658.2005.04864.x

Chen Y, Du H. The promising PARP inhibitors in ovarian cancer therapy: From Olaparib to others. Biomedicine & Pharmacotherapy. 2018 Mar 1; 99:552–60. 10.1016/j.biopha.2018.01.094

Maynard S, Schurman SH, Harboe C, de Souza-Pinto NC, Bohr VA. Base excision repair of oxidative DNA damage and association with cancer and aging. Carcinogenesis. 2009 Jan 1;30(1):2–10. 10.1093/carcin/bgn250

Kleine H, Poreba E, Lesniewicz K, Hassa PO, Hottiger MO, Litchfield DW, Shilton BH, Lüscher B. Substrate-assisted catalysis by PARP10 limits its activity to mono-ADP-ribosylation. Molecular cell. 2008 Oct 10;32(1):57–69. 10.1016/j.molcel.2008.08.009

Rojo F, Garcia-Parra J, Zazo S, Tusquets I, Ferrer-Lozano J, Menendez S, Eroles P, Chamizo C, Servitja S, Ramírez-Merino N, Lobo F. Nuclear PARP-1 protein overexpression is associated with poor overall survival in early breast cancer. Annals of oncology. 2012 May 1;23(5):1156–64. 10.1093/annonc/mdr361

Nilov DK, Pushkarev SV, Gushchina IV, Manasaryan GA, Kirsanov KI, Švedas VK. Modeling of the Enzyme—Substrate Complexes of Human Poly (ADP-Ribose) Polymerase 1. Biochemistry (Moscow). 2020 Jan; 85:99–107. 10.1134/S0006297920010095

Langelier MF, Pascal JM. PARP-1 mechanism for coupling DNA damage detection to poly (ADP-ribose) synthesis. Current opinion in structural biology. 2013 Feb 1;23(1):134–43. 10.1016/j.sbi.2013.01.003

LaFargue CJ, Dal Molin GZ, Sood AK, Coleman RL. Exploring and comparing adverse events between PARP inhibitors. The Lancet Oncology. 2019 Jan 1;20(1): e15–28. 10.1016/S1470-2045(18)30786-1

Mughal, T. A., Ali, S., Mumtaz, S., Summer, M., Saleem, M. Z., Hassan, A., & Hameed, M. U. (2024). Evaluating the biological (antidiabetic) potential of TEM, FTIR, XRD, and UV-spectra observed berberis lyceum conjugated silver nanoparticles. Microscopy Research and Technique, 87(6), 1286–1305.

Tahir, H., Rashid, F., Ali, S., Summer, M., & Abaidullah, R. (2024). Spectrophotometrically, spectroscopically, microscopically and thermogravimetrically optimized TiO2 and ZnO nanoparticles and their bactericidal, antioxidant and cytotoxic potential: a novel comparative approach. Journal of Fluorescence, 34(5), 2019–2033.

Tanveer, T., Ali, S., Ali, N. M., Farooq, M. A., Summer, M., Hassan, A., … & Islam, R. (2024). Evaluating the effect of pH, temperature and concentration on antioxidant and antibacterial potential of spectroscopically, spectrophotometrically and microscopically characterized Mentha Spicata capped silver nanoparticles. Journal of Fluorescence, 34(3), 1253–1267.

Akhtar, M. F., Irshad, M., Ali, S., Summer, M., Gulrukh, S., & Irfan, M. (2024a). Evaluation of biological potential of UV-spectrophotometric, SEM, FTIR, and EDS observed Punica granatum and Plectranthus rugosus extract-coated silver nanoparticles: A comparative study. Microscopy Research and Technique, 87(3), 616–627.

Subhani, A. A., Irshad, M., Ali, S., Jawad, M., Akhtar, M. F., & Summer, M. (2024). UV-spectrophotometric optimization of temperature, pH, concentration and time for eucalyptus globulus capped silver nanoparticles synthesis, their characterization and evaluation of biological applications. Journal of Fluorescence, 34(2), 655–666.

Akhtar, M. F., Irshad, M., Ali, S., Summer, M., Jawad, M., Akhter, M. F., … & Asghar, G. (2024b). Spectrophotometric, microscopic, crystallographic and X-ray based optimization and biological applications of Olea paniculata leaf extract mediated silver nanoparticles. South African Journal of Botany, 166, 97–105.

Li, L., Tan, J., Miao, Y., Lei, P., & Zhang, Q. (2015). ROS and autophagy: interactions and molecular regulatory mechanisms. Cellular and molecular neurobiology, 35, 615–621. 10.1007/s10571-015-0166-x

Collins, Y., Chouchani, E. T., James, A. M., Menger, K. E., Cochemé, H. M., & Murphy, M. P. (2012). Mitochondrial redox signalling at a glance. Journal of cell science, 125(4), 801–806. 10.1242/jcs.098475

Agarwal S, Mehrotra RJ. An overview of molecular docking. JSM chem. 2016;4(2):1024–8. ISSN: 2333-6633

Cortez AJ, Tudrej P, Kujawa KA, Lisowska KM. Advances in ovarian cancer therapy. Cancer chemotherapy and pharmacology. 2018 Jan; 81:17–38. 10.1007/s00280-017-3501-8

Adnan M, Siddiqui AJ, Hamadou WS, Patel M, Ashraf SA, Jamal A, Awadelkareem AM, Sachidanandan M, Snoussi M, De Feo V. Phytochemistry, bioactivities, pharmacokinetics and toxicity prediction of Selaginella repanda with its anticancer potential against human lung, breast and colorectal carcinoma cell lines. Molecules. 2021 Feb 2;26(3):768. 10.3390/molecules26030768

Kunert O, Swamy RC, Kaiser M, Presser A, Buzzi S, Rao AA, Schühly W. Antiplasmodial and leishmanicidal activity of biflavonoids from Indian Selaginella bryopteris. Phytochemistry letters. 2008 Dec 12;1(4):171–4. 10.1016/j.phytol.2008.09.003

Shridhar Deshpande N, Mahendra GS, Aggarwal NN, Gatphoh BF, Revanasiddappa BC. Insilico design, ADMET screening, MM-GBSA binding free energy of novel 1, 3, 4 oxadiazoles linked Schiff bases as PARP-1 inhibitors targeting breast cancer. Future Journal of Pharmaceutical Sciences. 2021 Aug 28;7(1):174. 10.1186/s43094-021-00321-4

Muhammad Tahir, H., Saleem, F., Ali, S., Ain, Q. U., Fazal, A., Summer, M., … & Murtaza, G. (2020). Synthesis of sericin-conjugated silver nanoparticles and their potential antimicrobial activity. Journal of basic microbiology, 60(5), 458–467.

Summer, M., Tahir, H. M., Ali, S., Abaidullah, R., Mumtaz, S., Nawaz, S., & Azizullah. (2023). Bactericidal potential of different size sericin-capped silver nanoparticles synthesized by heat, light, and sonication. Journal of Basic Microbiology, 63(9), 1016–1029.

Mumtaz, S., Ali, S., Kazmi, S. A. R., Mughal, T. A., Mumtaz, S., Tahir, H. M., … & Rashid, M. I. (2023). Analysis of the antimicrobial potential of sericin-coated silver nanoparticles against human pathogens. Microscopy Research and Technique, 86(3), 320–330.

Hanwell, M. D., Curtis, D. E., Lonie, D. C., Vandermeersch, T., Zurek, E., & Hutchison, G. R. (2012). Avogadro: an advanced semantic chemical editor, visualization, and analysis platform. Journal of cheminformatics, 4, 1–17. 10.1186/1758-2946-4-17

Morris GM, Huey R, Lindstrom W, Sanner MF, Belew RK, Goodsell DS, Olson AJ. AutoDock4 and AutoDockTools4: Automated docking with selective receptor flexibility. Journal of computational chemistry. 2009 Dec;30(16):2785–91. 10.1002/jcc.21256

Tian, W., Chen, C., Lei, X., Zhao, J., & Liang, J. (2018). CASTp 3.0: computed atlas of surface topography of proteins. Nucleic acids research, 46(W1), W363–W367. 10.1093/nar/gky473

Wadhwa, K., Kaur, H., Kapoor, N., & Brogi, S. (2023b). Identification of sesamin from sesamum indicum as a potent antifungal agent using an integrated in silico and biological screening platform. Molecules, 28(12), 4658. 10.3390/molecules28124658

Daina A, Michielin O, Zoete V. SwissADME: a free web tool to evaluate pharmacokinetics, drug-likeness and medicinal chemistry friendliness of small molecules. Scientific reports. 2017 Mar 3;7(1):42717. 10.1038/srep42717

Denizot F, Lang R. Rapid colorimetric assay for cell growth and survival: modifications to the tetrazolium dye procedure giving improved sensitivity and reliability. Journal of immunological methods. 1986 May 22;89(2):271–7. 10.1016/0022-1759(86)90368-6.

